# mTORC1-mediated acquisition of reward-related spatial representations by hippocampal somatostatin interneuronsa

**DOI:** 10.1101/2023.02.24.529922

**Authors:** François-Xavier Michon, Isabel Laplante, Anthony Bosson, Richard Robitaille, Jean-Claude Lacaille

## Abstract

Plasticity of principal cells and inhibitory interneurons underlies hippocampal memory. Bidirectional modulation of somatostatin cell mTORC1 activity, a crucial translational control mechanism in synaptic plasticity, causes parallel changes in hippocampal CA1 somatostatin interneuron (SOM-IN) long-term potentiation and hippocampus-dependent memory, indicating a key role in learning. However, SOM-IN activity changes and behavioral correlates during learning, and the role of mTORC1 in these processes, remain ill-defined. To address these questions, we used two-photon Ca^2+^ imaging from SOM-INs during a virtual reality goal-directed spatial memory task in head-fixed control mice (SOM-IRES-Cre mice) or in mice with conditional knockout of *Rptor* (SOM-Rptor-KO mice) to block mTORC1 activity in SOM-INs. We found that control mice learn the task, but SOM-Raptor-KO mice exhibit a deficit. Also, SOM-IN Ca^2+^ activity became increasingly related to reward localization during learning in control mice but not in SOM-Rptor-KO mice. Four types of SOM-IN activity patterns related to reward location were observed, “reward off sustained”, “reward off transient”, “reward on sustained” and “reward on transient”, and these responses showed global remapping after reward relocation in control but not SOM-Rptor-KO mice. Thus, SOM-INs develop mTORC1-dependent spatial coding related to learning reward localization. This coding may bi-directionally interact with pyramidal cells and other structures to represent and consolidate reward location.

## Introduction

Learning and memory are essential functions for animal survival that involves neuronal networks in several regions and cellular mechanisms such as long-term synaptic plasticity. Regulation of excitatory synaptic transmission and activity of CA1 pyramidal cells (PCs) is under strong inhibitory control by feedforward and feedback inhibitory circuits (1–4). These complex inhibitory interconnections contribute to the modulation of hippocampal networks, as well as to the formation and coordination of neuronal assemblies underlying learning and memory (3, 5–7).

Hippocampal inhibitory interneurons (INs) are heterogeneous populations of GABAergic inhibitory cells with varied morphological, molecular, and electrophysiological properties, as well as specialized network functions (7–14). In CA1 hippocampus, somatostatin-expressing interneurons (SOM-INs) are a major subgroup of INs which include Oriens-Lacunosum/Moleculare (O-LM) cells, bistratified cells and long-range projecting cells (4, 15, 16). SOM-INs have a key role in regulation of PC activity, as well as hippocampal learning and memory (5, 16–19). SOM-INs modulate the spiking rate and burst firing of PCs in vitro (3) and reduce the activity of place cells during exploration in vivo (5). Furthermore, silencing SOM-INs during learning impairs long-term contextual memory (17, 19) and object location memory (20). More recently, the coupling of in vivo calcium imaging and immunohistochemical identification of CA1 IN subtypes in mouse during head-fixed exploration and goal-directed learning, showed preferential recruitment of IN subtypes with quantitative differences in response properties and feature selectivity during key behavioral tasks and states (21). For SOM-INs, their activity is tied to animal movement, as SOM-IN activity is correlated with animal locomotion and most cells increasing their activity during movement (22). In addition SOM-IN activity is modulated by spatial learning (21). Thus, SOM-INs could regulate memory formation depending on the brain state and action of the animal.

Long-term potentiation (LTP) of synapses is a main cellular mechanism involved in memory that has mainly been examined at excitatory synapses onto PCs (23). However, recent studies revealed that excitatory synapses onto SOM-INs also undergo LTP (16, 18, 24, 25). LTP in SOM-INs requires the activation of mGluR1a, a metabotropic glutamate receptor subunit highly expressed in SOM-INs and which triggers the synthesis of new proteins (18, 19, 26–28). A key protein complex involved in the regulation of protein synthesis during LTP through activation of mGluR1, is the mechanistic target of rapamycin complex 1 (mTORC1) (29). Bidirectional modulation of mTORC1 activity in SOM-INs causes parallel changes in learning-induced LTP in SOM-INs and in hippocampal memory. Cell-specific conditional knock-out in SOM-INs of the gene expressing the Regulatory-Associated Protein of mTOR (Raptor), which is a necessary component of mTORC1, reduces mTORC1 activity in SOM-INs, prevents mGluR1a-mediated LTP in SOM-INs, and causes impairment of long-term contextual fear and spatial memory (18). Conversely, cell-specific conditional heterozygous knock-out in SOM-INs of the Tuberous Sclerosis Complex 1 (TSC1) gene, a repressor of mTORC1, increases mTORC1 activity in SOM-INs, facilitates induction of mGluR1a-mediated LTP in SOM-INs, and results in facilitation of long-term contextual fear and spatial memory (18). Thus mTORC1-mediated synaptic plasticity in SOM-INs is crucial for hippocampal learning.

However, changes in SOM-IN activity, and their behavioral correlates, during learning of a spatial memory task remain to be determined. Moreover, the role of mTORC1 function and plasticity in SOM-IN firing changes during learning has not been examined. In the present study, we address these questions by combining in vivo 2-photon calcium imaging of SOM-INs and learning of a virtual reality goal-directed spatial memory task, in control mice expressing Cre in SOM-INs (SOM-IRES-Cre mice) and in mice with a conditional knockout of *Rptor* in SOM-INs (SOM-Rptor-KO mice) for cell-specific impairment of mTORC1 activity and plasticity. We found that control mice learned the virtual spatial memory task but that SOM-Raptor-KO mice showed a learning deficit. Moreover, SOM-INs of control mice acquired place activity related to reward location during learning. Four types of spatial activity were distinguished during learning: “reward off sustained”, “reward on sustained”, “reward off transient” and “reward on transient” responses. However, SOM-INs of SOM-Raptor-KO mice failed to acquire such reward-related place activity during learning. Furthermore, SOM-IN reward-related place activity in control mice was sensitive to reward relocation. Global remapping of SOM-IN activity occurred with relocation of the reward to a different area of the maze, but not in SOM-Raptor-KO mice. Our results show that SOM-INs acquire a mTORC1-dependent place activity during learning of a spatial memory task, indicating a major role of SOM-INs in representation of reward location and of mTORC1 in the acquisition of learning-related SOM-IN firing correlates. Moreover, the learning impairment in SOM-Raptor-KO mice was associated with a deficit in learning-related activity of SOM-INs. Thus, SOM-IN spatial activity is unlike the typical place field activity of PCs and may bi-directionally interact with PCs to represent and consolidate memory of reward location.

## Results

### Goal-directed spatial learning is impaired by conditional knock-out of Rptor in SOM-INs

To investigate the activity of SOM-INs, their behavioral correlates, and the role of mTORC1 during learning, we developed a virtual reality goal-directed spatial memory task in head-fixed control mice (*Sst*^ires-Cre/wt^ mice; called SOM-IRES-Cre mice) (Fig. 1) and in mice with a conditional knockout of *Rptor* in SOM-INs (*Sst*^ires-Cre/wt^;*Rptor*^fl/fl^ mice; called SOM-Rptor-KO mice).

**Fig. 1.**
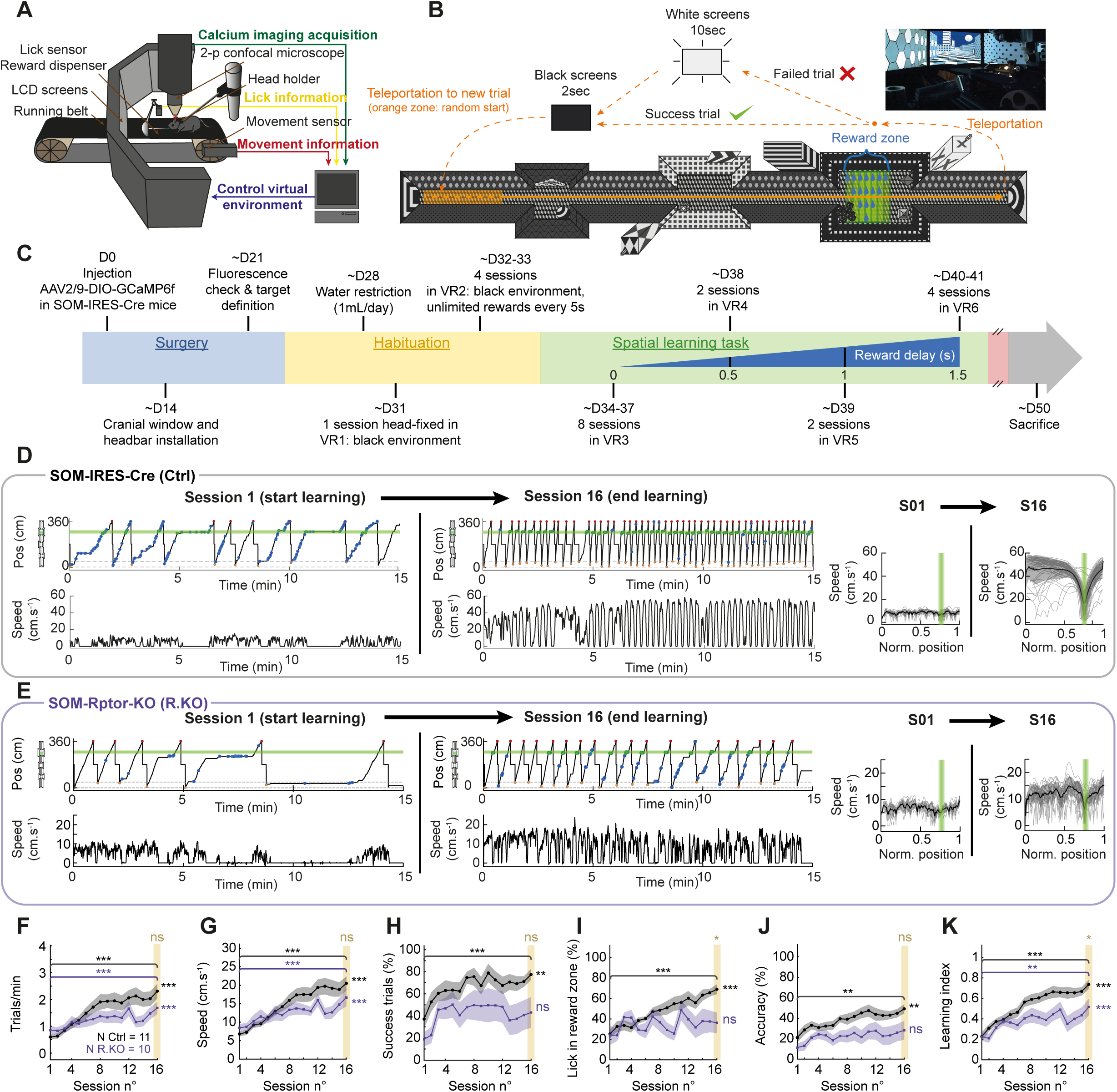
Learning of goal-directed spatial memory task in control mice and deficit in SOM-Rptor-KO mice. **A** Diagram of the virtual reality, treadmill, head-fixation, and 2-photon microscope setup. **B** Top view illustration of the virtual environment for the spatial learning task. **C** Timeline of surgical and behavioral procedures. **D** Left: Position and associated speed signals of a representative control SOM-IRES-Cre mouse during session 1 (start of learning) and session 16 (end of learning), showing an increase in number of trials per session, animal speed, and licks in the reward zone at the end of training. Green shading corresponds to reward area. Blue circle/blue outline indicates lick outside the reward zone, green circle/green outline lick in the reward zone and reward, and green circle/blue outline lick in the reward zone but reward was no longer available. Right: Speed profile as function of position (mean of all trials in black, individual trials in grey) for sessions 1 and 16. Green shading corresponds to reward zone. **E** Similar representation of training of a representative SOM-Rptor-KO mouse, showing comparable increase in number of trials per session and animal speed with training, but with increased licks unrelated to reward. **F**, **G**, **H**, **I**, **J**, **K** Summary plots of changes over training sessions in SOM-IRES-Cre (n=11 mice; Ctrl; black) and SOM-Rptor-KO (n=10 mice; R.KO; magenta) mice showing similar increases in number of trials per minute (**F**) and animal speed (**G**) in both mice groups; increases in percent success trials (**H**), percent lick in reward zone (**I**), and lick accuracy (**J**) only in control SOM-IRES-Cre mice; and reduced learning index (**K**) in SOM-Rptor-KO mice relative to control SOM-IRES-Cre mice, indicative of a spatial learning deficit in SOM-Rptor-KO mice. Details of statistical tests provided in Additional file 5: Table S1. * p<0.05, ** p<0.01, *** p<0.001, ns not significant.

First, we verified that the single allele-driven expression of Cre-recombinase in SOM-INs and conditional homozygous knock-out of *Rptor* prevent mTORC1 signaling in SOM-INs, using an immunofluorescence assay of ribosomal protein S6 phosphorylation as previously (18, 24). In acute hippocampal slices from control mice, chemical induction of mGluR1- and mTORC1-mediated LTP (treatment with the mGluR1/5 agonist DHPG in presence of the mGluR5 antagonist MPEP) increased phosphorylation of S6 in EYFP-expressing SOM-INs relative to sham-treatment (Additional file 1: Fig. S1A-C). The same chemical induction protocol failed to induce S6 phosphorylation in SOM-INs of SOM-Rptor-KO mice (Additional file 1: Fig. S1C). Basal level of S6 phosphorylation was also reduced in EYFP-expressing SOM-INs of SOM-Rptor-KO mice relative to control mice (Additional file 1: Fig. S1B). These results indicate an efficient block of mTORC1 signaling in SOM-INs of SOM-Rptor-KO mice.

Next, we exposed SOM-IRES-Cre mice and SOM-Rptor-KO mice to the head-fixed virtual reality goal-directed learning task (Fig. 1A-C; see Methods for details). A first series of head-fixed experiments were performed with behavioral analysis only (without cranial window and Ca^2+^ imaging) with 4 control SOM-IRES-Cre mice and 5 SOM-Rptor-KO mice. Subsequent experiments combined behavior with Ca^2+^ imaging and were carried out with an additional 7 control SOM-IRES-Cre mice and 5 SOM-Rptor-KO mice. Behaviors were generally similar in mice with or without cranial windows, and results were pooled together for behavioral data analysis. Briefly, after a recovery period from surgery, mice were given a period of habituation of 5 daily sessions with the experimental set-up (head-fixed on running belt, no virtual reality maze, 4 sessions with non-specific reward) (Fig. 1C). Next mice were trained twice a day for 16 sessions in the goal-oriented spatial learning task (Fig. 1B). For each training trial, the mouse was teleported to a random location in a start zone of the linear maze that was projected on LCD screens. Movement of the running belt by the mouse translated in movement in the virtual maze. The animal was trained to obtain reward (sweetened liquid) after stopping and licking at a specific location in the virtual maze. After continuing to the end of the maze, the animal was teleported back to the start zone for another trial. The behavior of the animal was quantified in terms of position in the maze, movement (belt speed), licking and reward.

Control mice learned the task gradually over the training sessions (Fig. 1D, F-K). The behavior of a representative control mouse during session 1 (start learning) and session 16 (end learning) is illustrated in Fig. 1D. Over training, animals showed an increase in the number of trials per session and in movement speed (Fig. 1D), as well as a decrease in trial duration (Additional file 1: Fig. S1D). At first, mice tended to lick over the entire length of the maze. But with training, animals licked more inside the reward zone than outside (Fig. 1D, I). They also showed increases in successful trials (Fig. 1D, H), accuracy of licks in reward zone (Fig. 1J), and time in the reward zone (Additional file 1: Fig. S1E). The animal velocity profile during the task became increasingly stereotyped with animals running faster toward the reward, stopping at the reward zone, and resuming running to the end of the maze (Fig. 1D). To quantify learning in the reward-directed spatial memory task, we considered that learning occurred when the animal retrieved rapidly and accurately the reward. So we quantified the number of trials per minute, the overall speed of movement, the percentage of success trials, the percentage of licks in the reward zone, and the accuracy of the licks over all sessions of training (Fig. 1F-J), and scored these measures with the same weight to determine an overall learning index (between 0 and 1) that reflects learning of the task in control animals (Fig. 1K; see Methods for details).

In contrast to control mice, SOM-Rptor-KO mice showed an impairment in learning the spatial memory task (Fig. 1E, F-K). Similar to control mice, SOM-Rptor-KO mice showed an increase in trial per minutes (Fig. 1F) and animal speed (Fig. 1G) with training. However, they failed to learn the goal-directed spatial nature of the task, i.e. to run to the reward zone, stop and lick to obtain reward. SOM-Rptor-KO mice failed to show improvement over training in success trials, licks in reward zone, and accuracy of reward (Fig. 1H-J). So, although the learning index increased with training in SOM-Rptor-KO mice, it was impaired relative to control mice (Fig. 1K). These results indicate that mTORC1 function in SOM-INs is required for learning the goal-directed spatial memory task.

### SOM-IN activity related to reward location

Having established that control mice learn the goal-directed spatial memory task and that mTORC1 function in SOM-INs is required, next we investigated the activity of SOM-INs and the role of mTORC1 function during learning, using 2-photon Ca^2+^ imaging and expression of the genetically encoded Ca^2+^ indicator GCaMP6f in hippocampal CA1 SOM-INs of SOM-IRES-Cre and SOM-Rptor-KO mice (Fig. 2A).

**Fig. 2.**
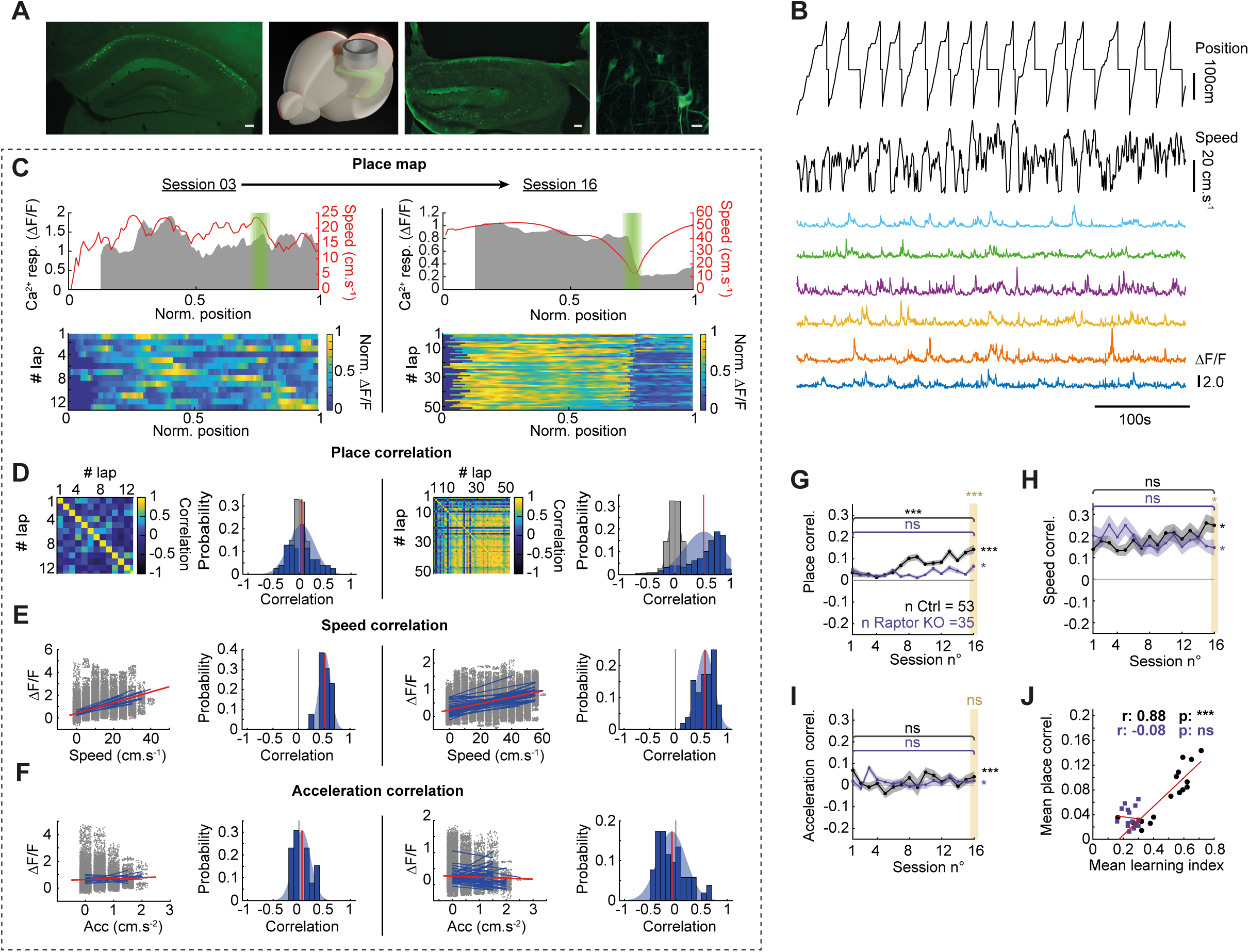
SOM-IN activity correlates with reward-related position in maze during training but not in SOM-Rptor-KO mice. **A** From left to right: (left most) fluorescence image of GCaMP6f expression in hippocampal SOM-INs of a representative control animal without cranial surgery; scale bar 100 µm. (Middle left) Diagram showing cranial window position above dorsal hippocampus. (Middle right) Fluorescent image from representative animal after behavior experiments showing neocortical tissue removed for cranial window placement and hippocampal SOM-IN GCaMP6f expression; scale bar 100 µm. (Rightmost) High power confocal images of in vivo field of view with SOM-INs expressing GCaMP6f; scale bar 20 µm. **B** Example of simultaneous measurements (from top to bottom) of maze position, animal speed, and Ca^2+^ responses (colored traces) from 6 SOM-INs during a training session. **C** Ca^2+^ responses of a representative SOM-IN at start (session 3; left) and end (session 16; right) of training. Top: mean Ca^2+^ responses (grey) and speed (red) as function of position for all trials with reward zone indicated in green. Bottom: color-coded Ca^2+^ activity in each trial of the session, showing activity correlated with position at the end of training. **D** Correlation of Ca^2+^ responses with position across laps (place correlation) at sessions 3 and 16, showing place correlation developed with training. For each left: place correlation matrix of all paired laps. For each right: distribution of r values (blue), mean r (red) versus r distribution obtained by shuffling position measures (gray). **E** Correlation of Ca^2+^ responses with animal speed is present at sessions 3 and 16. For each left: Ca^2+^ responses as function of speed, blue lines correspond to correlation for each trial and red line to mean. For each right: distribution of r values (blue), mean r (red). **F** Similar representation of Ca^2+^ responses correlation with animal acceleration that is absent at sessions 3 and 16. **G** Mean place correlation for all SOM-INs showing the increase in place correlation over training sessions in control mice but not in SOM-Rptor-KO mice. **H**, **I** Mean correlation of Ca^2+^ activity with speed (**H**) and acceleration (**I**) for all SOM-INs did not change with training. **J** Mean place correlation as a function of mean learning index over training sessions for all animals, showing correlation in control but not SOM-Rptor-KO mice. Details of statistical tests provided in Additional file 5: Table S1. * p<0.05, ** p<0.01, *** p<0.001, ns not significant.

SOM-INs in the stratum o*riens/alveus* region with Ca^2+^ activity recorded for at least ¾ of the training sessions were considered for analysis. Ca^2+^ signals were monitored in 53 cells in 7 SOM-IRES-Cre mice and 35 cells in 5 SOM-Rptor-KO mice (7.33 ± 2.93 cells per mouse) over training sessions of the virtual reality spatial memory task. Ca^2+^ responses (ΔF/F) of SOM-INs were measured during each session (Fig. 2B) and the following correlated behavioral variables were examined: position of the animal in the maze (Fig. 2C, D), animal speed (Fig. 2E), acceleration (Fig. 2F) and deceleration (Additional file 2: Fig. S2A). Ca^2+^ responses and behavioral correlates for the first and last training sessions are illustrated in Fig. 2C for a representative SOM-IN from a control SOM-IRES-Cre mouse. For this SOM-IN, Ca^2+^ activity was not related to the animal location in the maze at the start of training, but it was at the end of training. SOM-IN Ca^2+^ activity related to reward location was different from the place cell encoding of CA1 pyramidal neurons (30–32). In this SOM-IN example, Ca^2+^ activity did not increase or decrease as the animal passes through a specific location but was elevated as the animal moved to the reward zone, decreased rapidly at the reward location, and remained low until the end of the trial. Thus, instead of a classical place field analysis, we considered the similar variations in Ca^2+^ signal through several passages on the track as the most important feature of the place coding of SOM-INs. We computed for each cell and session, the place correlation matrix by determining the place activity correlation for all pairs of trials and an overall mean place correlation value, and we compared it to a random distribution obtained by shuffling Ca^2+^ signals (Fig. 2D). In addition, correlation of Ca^2+^ activity with animal speed, acceleration or deceleration were obtained by binning measures and fitting a linear correlation for each trial (Fig. 2E, F). For the SOM-IN shown in Fig. 2C, Ca^2+^ activity was not correlated to location at start of training (r = 0.041 ± 0.0285) but showed clear place correlation at the end of training (Fig. 2D; r = 0.5113 ± 0.0095). In contrast, the SOM-IN Ca^2+^ activity correlation with animal speed previously reported (21, 22) was present at the start of training in this SOM-IN and did not change with training (Fig. 2E; r = 0.5208 ± 0.0360 at start; r = 0.5786 ± 0.0234 at end). Ca^2+^ activity of this SOM-IN was not correlated with acceleration at the start of training, nor at the end (Fig. 2F; r = 0.0295 ± 0.0472 at start; r = -0.0774 ± 0.0386 at end).

To characterize the spatial coding of SOM-INs, we analyzed the Ca^2+^ activity correlations for each SOM-IN over training sessions from control SOM-IRES-Cre mice and SOM-Rptor-KO mice. Place correlation of SOM-INs improved gradually throughout training and was increased at the end relative to the start of training in control mice but not in SOM-Rptor-KO mice (Fig. 2G). Interestingly, correlation with animal speed was not different between start and end of training in control SOM-IRES-Cre and SOM-Rptor-KO mice (Fig. 2H). Ca^2+^ activity was not correlated with acceleration and the lack of correlation was similar at the start and end of training for both mouse genotypes (Fig. 2I). Finally, Ca^2+^ activity correlation with deceleration was increased at the end relative to the start of training in control mice but not in SOM-Rptor-KO mice (Additional file 2: Fig. S2A). Together these results indicate that SOM-INs acquire place coding during training in the goal-directed spatial memory task in control mice. The increase in place correlation is associated with an increase in deceleration correlation but with no changes in speed and acceleration correlations. Moreover, the acquisition of place coding by SOM-INs over training was deficient in SOM-Rptor-KO mice, suggesting it is dependent on intact mTORC1 function in SOM-INs.

The correlation features of SOM-IN activity were linked with behavioral performance. Importantly, place correlation of SOM-INs was well correlated with learning index over the training sessions in control mice (Fig. 2J). Despite a lack of change in speed correlation through training, speed correlation was well correlated with learning index (Additional file 2: Fig. S2B), as well as with place correlation (Additional file 2: Fig. S2E) in control mice. Acceleration correlation, however, was not correlated with learning index (Additional file 2: Fig. S2C). Deceleration correlation, another aspect of animal velocity that may involve different processes than speed or acceleration and that was negatively correlated with Ca^2+^ activity and decreased during training (Additional file 2: Fig. S2A), was also correlated with learning index (Additional file 2: Fig. S2D), suggesting a reduced influence of these processes in the modulation of SOM-INs activity across learning. Importantly, these correlations between SOM-IN activity and learning index were absent in SOM-INs of SOM-Rptor-KO mice (Fig. 2J; Additional file 2: Fig. S2B-E), suggesting a key role of SOM-IN mTORC1 function in the development of place coding and behavioral performance.

### Heterogeneity of SOM-IN activity related to reward location

Because most SOM-INs exhibited Ca^2+^ activity related to reward location during training, we characterized in more detail Ca^2+^ responses in relation to reward location. For each cell in every session, we compared Ca^2+^ activity in three parts of the virtual environment, before the reward zone, in the reward zone, and after the reward zone, and considered a reward location-related modulation when Ca^2+^ activity changed by at least 30% in or after the reward zone, relative to activity before the reward zone, (Fig. 3A, B).

**Fig. 3.**
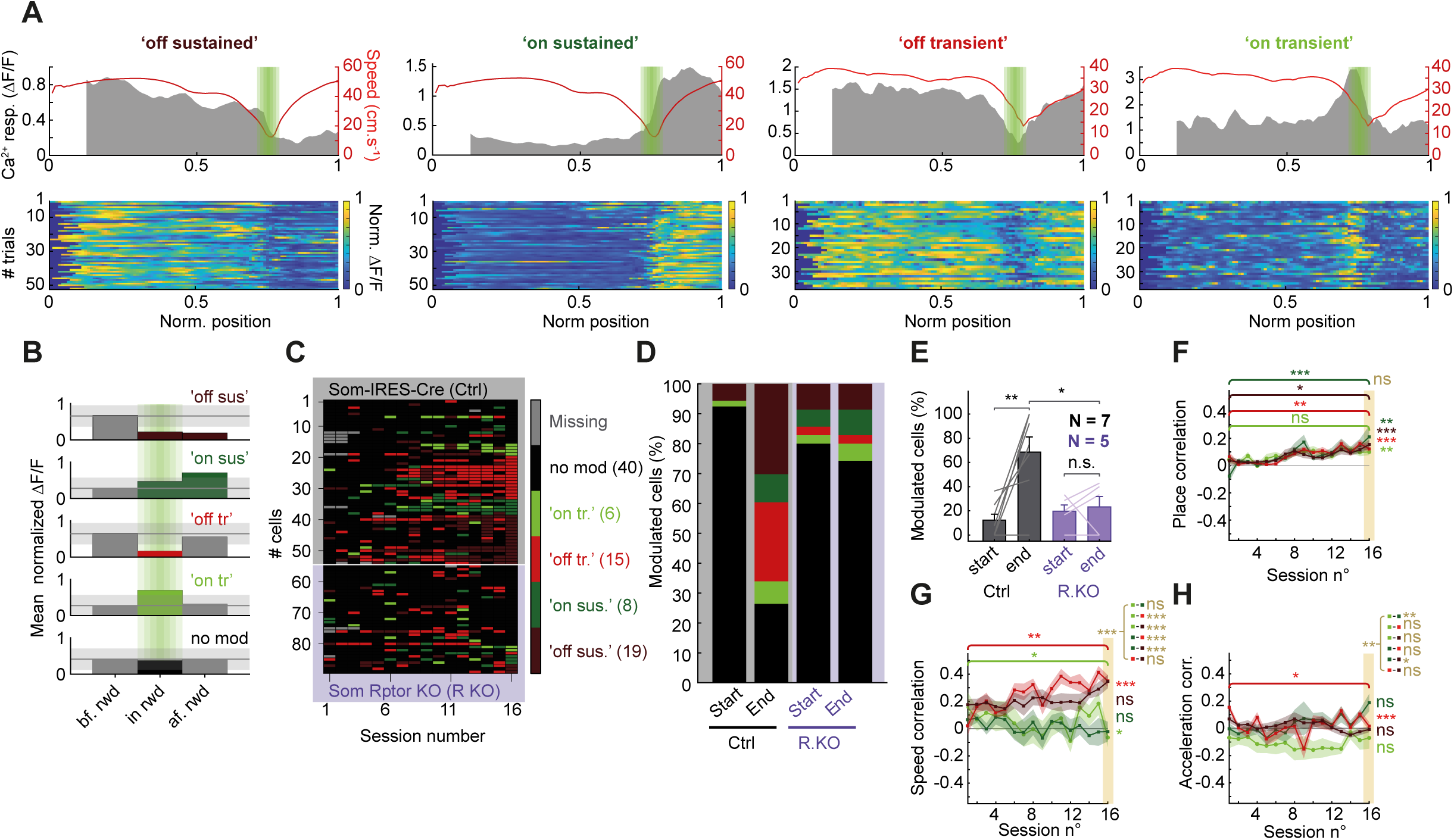
Heterogeneous SOM-IN Ca^2+^ activity related to reward location. **A** Representative examples of SOM-IN responses that developed during training: “off sustained”, “on sustained”, “off transient” and “on transient” responses. Top: mean Ca^2+^ responses (grey) and speed (red) as function of position for all trials in a session with reward zone indicated in green. Bottom: color-coded Ca^2+^ activity in each trial of the session. **B** Diagram of Ca^2+^ activity analysis criteria for each type of response with >30% changes (light gray shaded area) in Ca^2+^ activity in or after the reward zone, relative to activity before the reward zone (black line), defining response type (“off sustained”, dark red; “on sustained”, dark green; “off transient”, light red; “on transient”, light green; or no modulation, black). Representative examples of SOM-IN responses showing no modulation are given in Additional file 3: Fig. S3A. **C** Cell response identity matrix for all cells over training sessions ordered by response type at end of training, showing a gradual acquisition of spatial coding related to reward location. Top of matrix (grey): SOM-INs (n=53) from control mice. Bottom (magenta): SOM-INs (n=35) from SOM-Rptor-KO mice. **D** Distribution of cells with different response types at start and end of training for control and SOM-Rptor-KO mice, showing presence of 4 response types in both mouse genotypes, but increases with training in number of cells with responses only in control mice. **E** Summary graph for all cells with response related to reward location at start and end of training, showing increase in SOM-INs with responses in control (n=7) but not in SOM-Rptor-KO (n=5) mice. **F** Mean place correlation for SOM-INs with different response types (“off sustained” n =19; “on sustained” n =8; “off transient” n =15; “on transient” n =6), showing increase in place correlation over training for all response types. **G**, **H** Similar representation of Ca^2+^ activity correlation with speed (**G**) and acceleration (**H**) for SOM-INs with different response types, showing only few changes specific to certain response types. Details of statistical tests provided in Additional file 5: Table S1. * p<0.05, ** p<0.01, *** p<0.001, ns not significant.

With such criteria, four subtypes of Ca^2+^ responses were identified related to reward location (Fig. 3A-C). The most common response type (also shown in Fig. 2) consisted of elevated Ca^2+^ activity in the initial portions, followed by a decrease in activity in the reward zone which was maintained when the animal resumed movement after the reward. This type is referred to “reward off sustained” response (Fig. 3A) and they represented 30% of total SOM-INs at the end of learning in control mice (Fig. 3C, D). A second type, termed “reward on sustained’ response, was the opposite and consisted of low activity in the initial portions, followed by an increase in activity in the reward zone which was maintained after the reward zone, and it represented 9% of total SOM-INs in control mice (Fig. 3A, C-D). The other two types of response exhibited transient modulation in the reward zone, with the “reward off transient” type showing reduced activity and representing 26% of control SOM-INs, and the “reward on transient” type displaying increased activity and consisting of 8% of control SOM-INs (Fig. 3A, C-D). Other cells showed no change in activity in relation to the reward zone and represented 26% of SOM-INs at end of learning in control mice (Fig. 3C-D; Additional file 3: Fig. S3A).

When examining every cell response over training sessions, the reward location-related responses of SOM-INs of control mice appeared progressively during learning with less than 10% of cells showing response modulation at the start of training and 77% of cells showing responses at the end of learning (Fig. 3C-E). In addition, toward the end of training most SOM-INs tended to maintain the same response type from session to session (Fig. 3C).

In contrast, a few SOM-INs from SOM-Rptor-KO mice showed one of the four types of Ca^2+^ response related to reward location, but most cells did not (Fig. 3C). At the end of training, less SOM-INs of SOM-Rptor-KO mice (24%) showed reward location-related responses compared to control mice (68%; Fig. 3E). Moreover, the number of SOM-INs from SOM-Rptor-KO mice with responses related to reward location did not change over training (Fig. 3E). Furthermore, SOM-INs from SOM-Rptor-KO mice tended not to show the same response consistency toward the end of training as control SOM-INs (Fig. 3C). Together these results indicate that the acquisition of reward location-related place coding in SOM-INs during training requires mTORC1 function in SOM-INs.

Next, we examined if SOM-INs characterized by the different responses related to reward location, display similar Ca^2+^ activity correlation with behavioral variables during the task, as determined previously (Fig. 2). Interestingly, cells with any of the four types of response showed a similar increase in place correlation across training sessions (Fig. 3F), indicating consistent acquisition of the four different spatial coding responses over training. However, SOM-INs with different responses showed distinct relationship of Ca^2+^ activity with animal speed, acceleration, and deceleration (Fig. 3G, H; Additional file 3: Fig. S3B). SOM-INs with “reward off transient” responses showed an increase of speed correlation over training, whereas SOM-INs with “reward on transient” responses displayed a decrease in speed correlation (Fig. 3G), consistent with mice learning to stop in the reward zone. Despite the very low correlation with acceleration, SOM-INs with “reward off transient” responses showed a decrease in acceleration correlation over sessions (Fig. 3H). Correlation with deceleration was more complex (Additional file 3: Fig. S3B). SOM-INs with “reward on sustained”, “reward off transient” and “reward off sustained” responses increased deceleration correlation over training. However, SOM-INs with “reward on transient” responses increased anticorrelation with deceleration. Overall, these analyses show the acquisition of a strong relation of all types of SOM-IN activity with reward location in the maze as indicated by place correlation increases during learning, and a more complex and response-specific association with animal speed, acceleration, and deceleration in the maze. These activity correlations are linked due to the specific requirement of the task for the animal to stop at the reward location.

We also asked whether the correlation of activity of individual SOM-INs with each other during the task, which reflect the presence of different response types in the same animal, was different between control and SOM-Rptor-KO mice. The activity of SOM-INs in control mice showed less correlation with each other than in SOM-Rptor-KO mice (Additional file 3: Fig. S3C), suggesting that SOM-INs which lack mTORC1-mediated acquisition of spatial coding during learning, process information in a more homogeneous manner than control SOM-INs and lack the diversity of information processing required for learning the reward location-related spatial task.

### Remapping of SOM-IN activity with reward relocation

SOM-INs encompass multiple subtypes of interneurons in CA1 hippocampus (4, 16, 33). Thus, the heterogeneity of SOM-IN responses in the spatial memory task may be due to sampling different SOM-IN subtypes. Alternatively, a given SOM-IN may display different responses depending on the environment or context, a concept called remapping in hippocampal place cells (34). To investigate this question, control and SOM-Rptor-KO mice were exposed, after completion of the learning task, to a relearning task in the same environment but with the reward moved to a new location, and we examined the changes in SOM-IN activity (Fig. 4A).

**Fig. 4.**
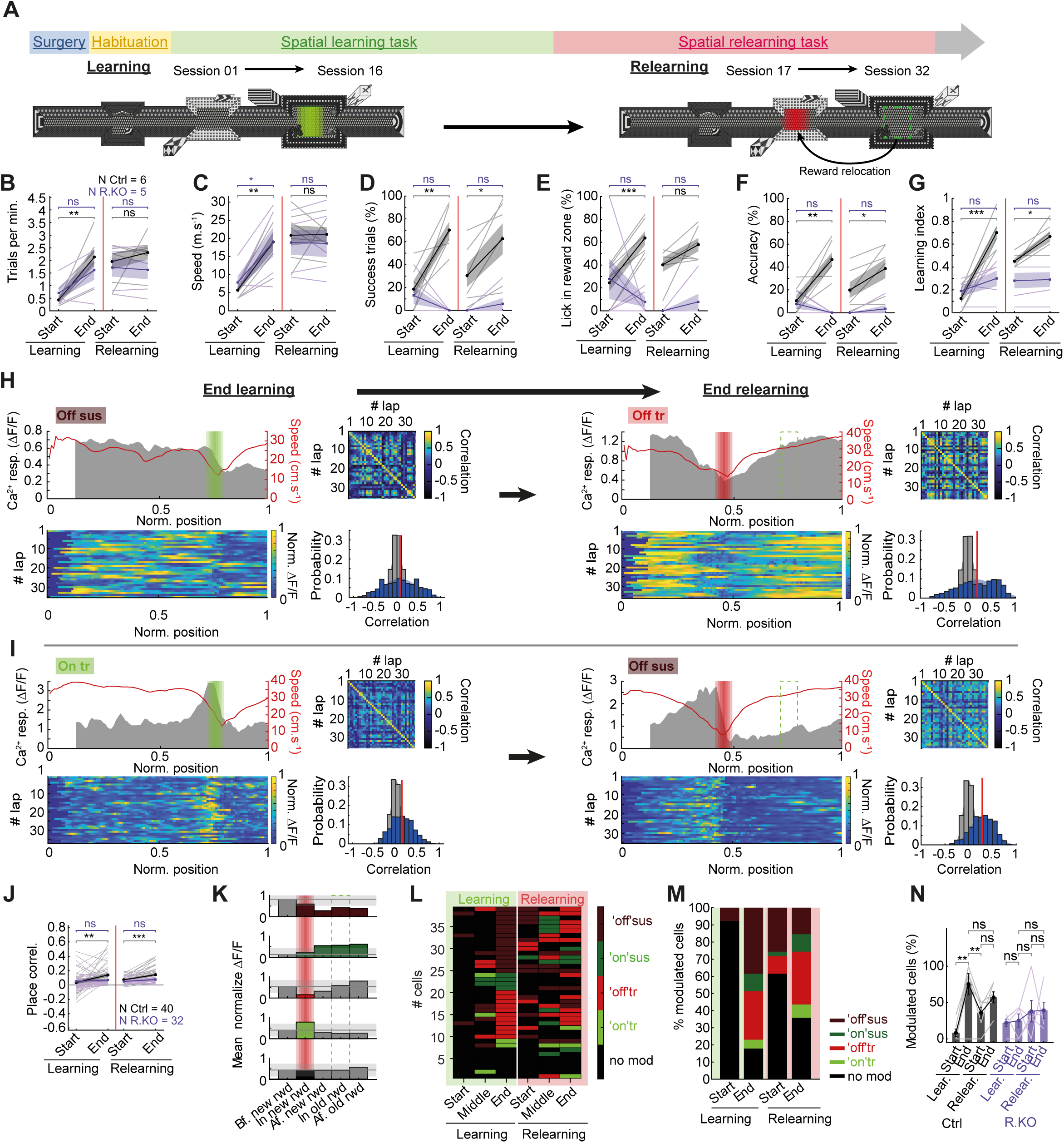
Remapping of SOM-IN activity with reward relocation. **A** Temporal sequence of surgical and behavioral procedures for relearning task with top view illustration of the virtual environment for the spatial learning task (reward location in green) and relearning task (new reward location in red; old location green dashed line). **B, C, D, E, F, G** Summary plots of behavioral changes during learning and relearning in control mice (n=6; black) and SOM-Rptor-KO mice (n=5; magenta) showing no changes in number of trials per minute (**B**), animal speed (**C**), and percent licks in reward zone (**E**) in both mice groups during relearning; increases in percent success trials (**D**), lick accuracy (**F**), and learning index (**G**) during relearning only in control SOM-IRES-Cre mice, showing that control mice, but not SOM-Rptor-KO mice, relearn a new reward location in the same environment. **H** Example of remapping of SOM-IN responses for a cell with “reward off sustained” response at end of learning (left) and “reward off transient” response at end of relearning (right). For each session, top left is mean Ca^2+^ responses (grey) and speed (red) as function of position for all trials in the session with reward zone indicated (green for learning; red for relearning); bottom left is color-coded Ca^2+^ activity in each trial of the session; top right is place correlation matrix of all paired laps; and bottom right is distribution of r values (blue), mean r (red) versus r distribution obtained by shuffling position measures (gray). **I** Similar representation of remapping for a SOM-IN with “reward on sustained” response at end of learning and “reward off sustained” response at end of relearning. **J** Mean place correlation with activity for all SOM-INs showing the increase during relearning in control but not SOM-Rptor-KO mice. **K** Diagram of Ca^2+^ activity analysis criteria for each type of response with >30% changes (light gray shaded area) in Ca^2+^ activity in the new reward zone (red shading) or after it, relative to activity before the reward zone (black line), defining response types (“off sustained”, dark red; “on sustained”, dark green; “off transient”, light red; “on transient”, light green; or no modulation, black) in the relearning task. **L** Cell response identity matrix for all cells in control mice during learning (green, left) and relearning (red, right) ordered by response type at end of learning, showing a gradual acquisition of a new spatial coding related to reward relocation during relearning. **M** Distribution of cells with different response types at start and end of learning and relearning for control mice, showing a decrease in number of modulated cells at start of relearning relative to end of learning, and an increase during relearning. **N** Summary graph for all cells with response related to reward location at start and end of learning and relearning, showing initial decrease and subsequent increase in SOM-INs with responses during relearning in control (n=6; black) but not in SOM-Rptor-KO (n=6; magenta) mice. Details of statistical tests provided in Additional file 5: Table S1. * p<0.05, ** p<0.01, *** p<0.001, ns not significant.

First at the behavioral level, the number of trials per minute and animal speed did not change at the start of relearning relative to the end of learning, and from the start to the end of relearning, in both control and SOM-Rptor-KO mice (Fig. 4B, C), consistent with mice having already learned during the learning phase to move quickly in the virtual environment to solve the task. In contrast, the performance measures related to learning the reward location (success trials, lick in reward zone, lick accuracy) decreased at the start of relearning relative to the end of learning, and then increased from start to end of relearning, in control mice (Fig. 4D-F), except for licks in reward zone that did not increase during relearning suggesting mice had already learned this aspect of the task. As a result, the learning index decreased at the start of relearning and increased from start to end of relearning in control mice (Fig. 4G), showing that mice relearn a new reward location in an unchanged environment. In contrast, SOM-Rptor-KO mice did not show changes in learning index and other measures related to learning reward location, at the start of relearning relative to the end of learning, and from start to end of relearning, indicating a deficit of relearning in these mice (Fig. 4D-G), and, thus, a requirement of intact mTORC1 function in SOM-INs for relearning in the spatial memory task. At the level of cell activity, we found that SOM-IN activity developed in relation to the new location of the reward during relearning in SOM-INs of control mice (Fig. 4H, I, K, L). Importantly, SOM-IN activity showed an increase in place correlation from start to end of relearning, as was the case during learning (Fig. 4J). At the start of relearning, the number of SOM-INs with responses related to reward location was reduced relative to the end of learning but then increased during relearning in control mice (Fig. 4L-N). Interestingly, the type of SOM-IN responses related to reward location changed during relearning in most SOM-INs of control mice (Fig. 4H, I, K, L). Many different permutations of response changes were observed (Fig. 4L). Notably, SOM-IN responses changed from “reward off sustained” in the learning task to “reward off transient” after relearning (Fig. 4H), from “reward off transient” to “reward off sustained” (Additional file 4: Fig. S4C), from “reward on transient” to “reward off sustained” (Fig. 4I), or some cells did not change their response type (Additional file 4: Fig. S4A) or others lost or gained response modulation (Additional file 4: Fig. S4B, D). The four types of reward location-related activity at the end of learning were present in similar proportion at the end of relearning, despite a change in response type in most cells (Fig. 4K-M). Finally, during relearning the correlation of activity with animal speed and acceleration did not change, but deceleration correlation was reduced (Additional file 4: Fig. S4E-G). Together these results indicate that SOM-INs acquire a new spatial coding of activity related to the relocation of reward, indicative of global remapping (34) during relearning of the goal-directed spatial memory task.

In contrast, the activity of SOM-INs of SOM-Rptor-KO mice did not change during relearning. Place correlation of SOM-IN activity was unchanged during relearning in SOM-Rptor-KO mice (Fig. 4J). Correlation of SOM-IN activity with animal speed, acceleration and deceleration were also unaffected (Additional file 4: Fig. S4E-G). The percent of modulated cells did not change at the start of relearning relative to the end of learning, nor from the start to the end of relearning, for SOM-INs of SOM-Rptor-KO mice (Fig. 4N), however the proportion of SOM-INs with “reward on sustained” responses increased (from 6% to 22%) with relearning in these mice (Additional file 4: Fig. S4H, I). These results indicate that global remapping of reward-related activity of SOM-IN during relearning is dependent on intact mTORC1 function in SOM-INs, suggesting a potential important functional role of mTORC1-dependent synaptic plasticity in SOM-INs in the process.

## Discussion

Our major findings are (1) control mice learn a virtual reality goal-directed spatial memory task but mice with impaired mTORC1 function in SOM-INs show a learning deficit; (2) Ca^2+^ activity of SOM-INs of control mice becomes related to reward localization during learning of the task, but not in SOM-INs with impaired mTORC1 function; and (3) four types of activity patterns related to reward location were observed SOM-INs, “reward off sustained”, “reward off transient”, “reward on sustained” and “reward on transient”, and these SOM-IN responses showed global remapping after relocation of the reward in control mice but not in mice with impaired mTORC1 function in SOM-INs. Thus, SOM-INs acquire, via mTORC1-dependent mechanisms, reward-related spatial coding activity that supports goal-directed spatial learning. The SOM-IN spatial activity related to reward location indicates a more complex modulation by locomotion and spatial learning than previously considered (21, 22), which may bi-directionally interact with the place field activity of hippocampal PCs to contribute to the hippocampal network representation and consolidation of reward location memory.

### mTORC1 mechanisms in SOM-INs

Our findings uncover a role of mTORC1 in the modification of Ca^2+^ activity of SOM-INs during spatial learning. It is noteworthy that mTORC1 function in SOM-INs does not appear essential for SOM-INs to display task-related spatial coding as mice with impaired mTORC1 function in SOM-INs learn the task and SOM-INs in these mice show some reward-related activity. So, mTORC1-independent mechanisms support a reduced level of SOM-IN activity changes and learning of the task. However, mTORC1 function is required for training-induced increases in reward-related SOM-IN responses and reward-related spatial learning of mice, indicating these mTORC1-mediated changes contribute to SOM-IN activity changes during learning and to spatial learning.

What mTORC1 mechanism might be involved? mTORC1 plays a central role in cell growth and metabolism (29). In mature neurons, mTORC1 is a key regulator of translation in long-term synaptic plasticity of principal cells and memory consolidation (35). In SOM-INs, mTORC1 plays an essential role in the control of protein synthesis during mGluR1a-mediated long-term potentiation (LTP) at pyramidal cell to SOM-IN (PC-SOM) synapses (18, 28). mTORC1-mediated PC-SOM synapse LTP, in turn, regulates CA1 hippocampal network metaplasticity, enhancing LTP at CA3 inputs and suppressing LTP at entorhinal inputs (6, 18, 19, 36). Moreover, mTORC1-mediated PC-SOM synapse LTP contributes to consolidation of long-term hippocampal spatial and contextual fear memory (18–20). Thus, an interesting possibility is that mTORC1-mediated plasticity at synapses of SOM-INs may be responsible for the changes in Ca^2+^ responses of SOM-INs that underlie place coding activity during training. Such a role of mTORC1 in translational control of synaptic plasticity in SOM-IN is consistent with previous work showing that eIF2a-mediated up-regulation of protein synthesis in hippocampal CA1 SOM-INs is sufficient to control hippocampal dependent long-term contextual fear memory (36). Thus, our results suggest that mGluR1a- and mTORC1-mediated LTP of SOM-INs may contribute to the acquisition of a reward location representation by SOM-INs. Our results provide new insight into the role in hippocampal memory function of long-term synaptic plasticity at excitatory synapses onto inhibitory interneurons (16, 37–39).

Other mTORC1 mechanisms may also be implicated in the changes in SOM-IN activity during learning of the spatial memory task. In trace eye-blink conditioning, an increase in intrinsic excitability of hippocampal CA1 SOM-INs, caused by a reduction in SK channels mediating slow afterhyperpolarizations, is associated with learning (40). However, whether mTORC1 is implicated in such changes during spatial learning is unknown. Moreover, mTORC1-mediated long-term potentiation of intrinsic excitability via downregulation of Kv1 channels occurs in CA1 parvalbumin interneurons but not in SOM-INs (41, 42). Also, axonal sprouting of SOM-INs and gain of dendritic inhibition associated with loss of parvalbumin synaptic inhibition in the CACNA1A mouse model of generalized epilepsy is reversed by rapamycin treatment and thus mTORC1-dependent (43). But how such axonal changes downstream of SOM-IN firing could be involved in spatial learning remains to be determined. Finally, in our study the conditional knockout of *Rptor* was global; thus, further approaches will be necessary to determine region-specific contributions of mTORC1 function in SOM-INs during spatial learning.

### Behavioral correlates of SOM-IN Ca^2+^ activity

At the cellular level, we found that the Ca^2+^ activity of SOM-INs of both genotypes were correlated with the animal speed during the task, and that speed correlation was unchanged at the end *versus* start of training (Fig. 2). The observed speed correlation is consistent with previous reports that SOM-IN activity during track running is primarily correlated with locomotion (21, 22) and occasionally with immobility (22).

It is important to note that the activity correlations with animal speed, acceleration, deceleration, and place location are linked because of the specific requirement of the task for the animal to stop at the reward location. Thus, velocity of the animal is intrinsically linked to the resolution of the task. A well-trained animal will stop at the reward location, thus intertwining the relation of activity with speed, acceleration, deceleration, and position in the maze. Thus, for cells with ‘reward on transient’ and ‘reward off transient’ responses, it is difficult to separate the influences of speed and reward location. This could also explain the observed correlation between velocity correlation and position correlation in control mice. However, correlation of activity with place increases with training but correlation with speed does not (Fig. 2), suggesting that SOM-INs do acquire some place activity related to reward location during learning.

In contrast, SOM-INs with ‘reward off sustained’ or ‘reward on sustained’ responses cannot be explained by a simple relationship with animal speed, acceleration, and deceleration. These reward location-related responses must involve movement-independent mechanisms for SOM-INs to remain active or inactive when locomotion starts again, and the animal leaves the reward zone. Thus, SOM-IN activity is partly determined by reward location during reward-related spatial learning. To determine the specific role of animal velocity in SOM-IN activity will require a different spatial memory task.

### Heterogeneity of SOM-IN Ca^2+^ responses

Our main result that SOM-INs acquire over training four types of activity patterns related to reward location, “reward off sustained”, “reward off transient”, “reward on sustained” and “reward on transient” suggests a more complex repertoire of SOM-IN activity related to reward location than previously reported. Imaging Ca^2+^ activity from immunohistochemically identified SOM-INs during a goal-oriented spatial learning task that involve mice running on a treadmill for water reward at a specific location, indicated that SOM-IN activity increased in the location immediately preceding the reward zone (21). Such responses are analogous to the SOM-IN responses in our experiments from cells becoming inactive in the reward zone (“reward off transient” and “reward off sustained” responses). Our results clearly show that SOM-INs also display responses characterized by increased activity in the reward zone (“reward on transient” and “reward on sustained responses”). The dichotomy in “on” and “off” responses related to reward location is consistent with the two types of activity related to movement or immobility found in CA1 SOM-INs during a virtual reality spatial navigation task (22). Complex types of responses related to reward location were also reported from SOM neurons recorded in prefrontal cortex of mice performing a reward foraging task (44). Two types of SOM neurons were identified as narrow spike SOM neurons (NS-SOM) or wide spike SOM neurons (WS-SOM). The activity of NS-SOM neurons was suppressed when entering the reward zone whereas activity of WS-SOM neurons was either suppressed or increased (44). Thus, SOM neurons in prefrontal cortex display heterogeneous responses related to reward location, analogous to our observations here, and these heterogenous responses may in part be due to recording from different subtypes of SOM neurons (44).

In our experiments we targeted expression of the Ca^2+^ indicator to all dorsal hippocampal CA1 SOM-INs and these include multiple subtypes of SOM-INs, including oriens lacunosum-moleculare (OLM) cells, bistratified cells, and cells with both local and long-range projections (oriens-retrohippocampal projecting cells, double projecting cells, and back-projecting cells) (16). Thus, a possibility is that the different types of Ca^2+^ responses related to reward location may be originate from different subtypes of SOM-INs. Interestingly, the prevalence of the distinct responses was uneven. After training, approximately 55% of SOM-INs showed reduced activity at reward location (“reward off” responses), 20% increased activity (“reward on” responses), and 25% no modulation. Because we sampled SOM-INs located in stratum oriens of dorsal hippocampus, many of our recordings were likely from O-LM cells (4, 33). Since, O-LM cells have direct inputs from the septum known to be linked with theta rhythm and animal movement (3, 45, 46), SOM-INs with “reward off transient” responses likely included O-LM cells. Another subtype of hippocampal inhibitory interneuron that target other local interneurons and have long-range projections that could be implicated in “reward on” responses are the so-called TORO (theta-OFF/ripple ON) cells. The activity of TORO cells is negatively correlated with animal speed and locomotion, and some TORO cells were identified as SOM-INs (47). However, more definite evidence of cell type-specific SOM-IN responses related to reward location will require recording and manipulating activity from identified hippocampal SOM-IN subtypes in goal-oriented spatial memory tasks.

### Remapping of SOM-IN representations

Hippocampal place cell representations are known to remap in response to change in the environment (34). The activity of SOM-INs during spatial tasks is also known to change upon exposure to a novel context (21, 48). Similarly, we found that SOM-IN responses related to a reward zone remapped when the location of the reward zone was changed. Similar suppression of activity at the previous reward zone and increase in activity at the new reward zone was previously reported in SOM-INs performing a goal-oriented learning task (21). Our findings further indicate that remapping occurs for all four types of SOM-INs responses, and even in initially non-modulated cells. Moreover, our results show that SOM-IN remapping did not occur in mice with impaired mTORC1 function in SOM-INs, suggesting that remapping requires cell-autonomous mTORC1 mechanisms and do not simply reflect up- and down-stream network interactions.

Interestingly, the global remapping of SOM-IN responses upon relocation of the reward zone indicated that individual SOM-IN often display different responses during learning the first and relearning the second reward location (Fig. 4 and Additional file 4: Fig. S4). Our observations of different responses in individual cells after remapping suggest that the different types of responses were not due solely to sampling different subtypes of SOM-INs but to specific dynamic network activity dependent on reward location and driven by pyramidal cells and other inputs. The switching of responses during remapping suggests a large versatility for individual SOM-INs in encoding reward location and, thus, in their bilateral network interactions.

How could individual SOM-INs show different responses related to reward location? SOM-INs have complex interactions with the hippocampal network. SOM-INs receive major excitatory inputs from CA1 pyramidal cells (16) but also from septum (46). SOM-INs receive inhibition from local inhibitory interneurons mostly those expressing vasoactive intestinal peptide (VIP) (49) and from brainstem nucleus incertus (50). SOM-IN axons profusely contact pyramidal cell dendrites but also other local interneurons (46, 51) as well as long-range targets (16). Thus, the different SOM-IN responses may thus arise from dynamic interactions in this complex network. An interesting consideration is that responses of SOM-IN with increased *versus* decreased activity at reward location may involve dynamic interactions between inhibitory VIP interneurons and SOM-INs (49), as VIP interneurons activity is mostly positively, but in some cases negatively, correlated with animal velocity and reward location during spatial foraging and goal-oriented spatial learning tasks (52).

### Network interactions

During a spatial memory task, an animal’s environment is not represented uniformly in the hippocampus, as CA1 pyramidal cells have an over-representation, both quantitatively and qualitatively, of salient locations such as reward sites, relative to other maze sites (31, 53–58). Also, CA1 pyramidal cells do not function as independent coding units. Coordinated connectivity and plasticity between co-active pyramidal cells and associated inhibitory subnetworks allow selective responses initiated in individual cells to adapt to multicellular assemblies (59, 60). Our results suggest that that learning-induced mTORC1-mediated changes in synaptic plasticity of SOM-INs impact the reward-related spatial coding of SOM-INs and associated behaviors. We speculate that one of the roles of SOM-INs could be to regulate the activity of pyramidal neurons around the location of the reward to increase its salience and better represent the location of interest in memory. However, changes in place cell activity caused by disruption of mTORC1-mediated synaptic plasticity and spatial encoding of SOM-INs remains to be determined.

SOM-INs, specifically O-LM cells, project to the distal part of pyramidal cell dendrites in stratum lacunosum-moleculare which also receive excitatory inputs from entorhinal cortex projections to CA1 (4, 17, 33, 61, 62). As an animal crosses a place field, synaptic coupling of CA1 place cells is decreased with parvalbumin interneurons and increased with SOM-INs, causing a switch of pyramidal cell inhibition from perisomatic/proximal dendritic to distal dendritic compartments, and allowing CA3 excitatory inputs to gain control over entorhinal excitatory inputs in driving pyramidal cell firing (63). Therefore, mTORC1-mediated SOM-IN representations during spatial learning may contribute to spatial/contextual information encoding by CA1 pyramidal cells by promoting internal representations by the hippocampal CA3 pathway while dampening external representations via the extrahippocampal entorhinal inputs at longer time scales.

## Materials and methods

### Animals

Animal procedures and experiments were performed in accordance with the Université de Montréal Animal Care Committee regulations (Comité de Déontologie de l’Expérimentation sur les Animaux; CDEA Protocols # 19-003, 19-004, 20-001, 20-002, 21-001, 21-002, 22-008, 22-009) and the Canadian Council of Animal Care guidelines.

Knock-in mice with an internal ribosome entry site (IRES)-linked Cre recombinase gene downstream of the *Sst* locus (*Sst*^ires-Cre^ mice; The Jackson Laboratory, Bar Harbour, ME, JAX# 013044) were crossed with C57BL/6J mice to generate control *Rptor* wild-type mice (*Sst*^ires-Cre/wt^;*Rptor*^wt/wt^ mice; termed SOM-IRES-Cre mice). *Sst*^ires-Cre^ mice were crossed with floxed *Rptor* mice (The Jackson Laboratory, Bar Harbour, ME, JAX# 013188) for cell-specific knock-out of *Rptor* in SOM cells. Heterozygous offsprings were backcrossed with floxed *Rptor* mice to generate *Sst*^ires-Cre/wt^;*Rptor*^fl/fl^ mice (termed SOM-Rptor-KO mice). All strains were maintained on a C57BL/6J background. An initial subset of control experiments were performed with behavioral analysis only on four *Sst*^ires-Cre^;*Rosa26*^lsl-EYFP^ mice (Cre-dependent enhanced yellow fluorescent protein [EYFP] expression in SOM-INs) (18) and five SOM-Rptor-KO mice without cranial window. The experiments combining behavior and calcium imaging with implanted cranial windows were carried out on two *Sst*^ires-Cre^ mice, five SOM-IRES-Cre mice and five SOM-Raptor-KO mice. No behavioral difference was noted and all control animals (5 SOM-IRES-Cre mice, *2 Sst*^ires-Cre^ mice and 4 *Sst*^ires-Cre^; *Rosa26*^lsl-EYFP^ mice) were pooled for behavioral analysis. Mice were housed 2-4 animals per cage before the first surgery, and subsequently were housed singly. Mice were maintained on a 12 h light/dark cycle with all experimentation performed during the light phase. Food and water were provided ad libitum until after recovery from the second surgery, at which time mice were restricted to 1ml/day of water and their weight and health monitored daily.

### S6 immunophosphorylation assay

CA1 SOM-INs were identified using Cre-dependent EYFP expression. SOM-IRES-Cre and SOM-Rptor-KO mice (4-6 weeks) were given an intraperitoneal (IP) injection of ketamine (50 mg/kg) and xylazine (5 mg/kg) and placed in a stereotaxic frame (Stoelting, Wood Dale, IL). AAV2/9-EF1a-DIO-EYFP viral particles (0.8μl; 10^13^ GC/ml; Addgene #27056) were injected bilaterally into the CA1 hippocampus (coordinates relative to bregma: AP -2.46 mm, ML ±1.75 mm, and DV -1.5 mm) at a flow rate of 0.1 µL/min using a 10 μl Hamilton syringe. The needle was left in place for 5 min after injection. Immunofluorescence experiments were performed within 6-8 days after virus injection. Mice were deeply anesthetized with sodium pentobarbital (MTC Pharmaceuticals, Cambridge, Ontario, Canada) and perfused transcardially with ice-cold artificial cerebrospinal fluid (ACSF) containing (in mM): 110 choline-chloride, 2.5 KCl, 7 MgCl_2_, 26 NaHCO_3_, 7 dextrose, 1.3 ascorbic acid and 0.5 CaCl_2_, and saturated with 95% O_2_-5% CO_2_. Coronal hippocampal slices (300 µm thickness) were obtained using a vibratome (Leica VT 1000S, Germany) in sucrose enriched ice-cold oxygenated ACSF containing (in mM): 87 NaCl, 2.5 KCl, 2.5 NaHCO_3_, 0.5 CaCl_2_, 7 MgCl_2_, 1.25 NaHPO_4_, 25 dextrose and 75 sucrose. Slices were transferred into normal oxygenated ACSF at room temperature containing (in mM): 124 NaCl, 2.5 KCl, 1.25 NaH_2_PO_4_, 2 MgCl_2_, 2 CaCl_2_, 26 NaHCO_3_, 10 dextrose, 1.3 ascorbic acid. After 1 hour recovery period, slices received the chemical induction protocol for late LTP that consisted of three applications (10 min duration each at 30 min intervals) of the mGluR1/5 agonist (S)-3,5-dihydroxyphenylglycine (DHPG, 5 μM; Abcam) in the presence of the mGluR5 antagonist 2-methyl-6 (phenylethynyl)-pyridine (MPEP, 25 μM; Tocris Bioscience) in ACSF at 31-33°C as previously (18). Immediately after treatment, slices were fixed with 4% paraformaldehyde, cryoprotected 24 hours later in 30% sucrose, and re-sectioned using a freezing microtome (50 µm thickness; Leica SMR200R, Germany). Sections were permeabilized for 15 min with 0.3% Triton X-100 in phosphate-buffered saline (PBS) and unspecific binding was blocked with 10% normal goat serum in 0.1% Triton X-100/PBS (1h). Rabbit polyclonal phospho-S6 ribosomal protein (S240/244) (1/2000; Cell Signaling, Beverly, MA #5364, RRID:AB 10694233) was incubated 48 hours at 4°C. Sections were subsequently incubated at room temperature with Alexa 594-conjugated goat anti-rabbit IgGs (1/500; 90 min; Jackson Immunoresearch Laboratories, West Grove, PA). Images were acquired using a confocal microscope (LSM880; Carl Zeiss, Oberkochen, Germany) at excitation 488 and 543 nm. Images from different treatment/groups were acquired using the exact same parameters. Cell fluorescence was quantified using ImageJ software (National Institute of Health) by comparing integrated density in cells corrected for background. For each animal, 2-3 hippocampal slices received chemical induction and 2-6 sections from those slices were quantified. A total of 39-75 fields of view were analyzed per conditions (3-6 independent mice experiments).

### Viral injection and cranial window placement

A first surgery was performed for injection of GCaMP6f viral vector in CA1 hippocampus. Animals were anesthetized with an intraperitoneal injection of ketamine (100 mg/kg) and xylazine (10 mg/kg). A small amount of lidocaine (0.3mg/kg, 0.03mg/ml) was applied locally at the site of skin incision. A small hole (0.25mm diameter) was made in the skull above the unilateral virus injection site (relative to bregma: AP -2mm, ML -1.4mm, and DV -1.5mm). AAV2/9-EF1a-DIO-GCaMP6f viral particles (0.8µL; 1.8×10^13^ gc/ml; Centre de Neurophotonics, Université Laval, Québec, Canada) diluted at 1/10 in 5% glycerol PBS were injected with an automated micropump (Hamilton) at 0.1µL/min. After injection, the needle remained in place for 7 min, following which the skin incision was sutured. Animals received a subcutaneous injection of meloxicam (2.5mg/kg) and were given post-surgical care for two days.

After 14 days of recovery, a second surgery was performed to install a cranial window for calcium imaging and a titanium bar for head fixation. Animals were anesthetized as described above. A craniotomy (3mm diameter) was performed over the unilateral injection site and the cortex overlying the hippocampus was removed by aspiration. The top layers of the external capsule were removed, and the lower layers left intact. Aspiration was unilateral and removed part of the visual, somatosensory, and parietal cortices. Previous studies indicate that such surgery does not impair mouse behavior in numerous tasks, including virtual reality track running (64, 65). Although not tested in detail, no obvious behavioral deficit was observed in these mice compared to those with headplate only. A custom-made imaging cannula (outer diameter 3 mm, inner diameter 2.36 mm, height 1.5 mm; Canula (Microgroup, Medway, MA) with a 3mm round glass coverslip glued at the end, was fixed to the skull with dental cement (C&B Metabond, Parkell, Edgewood, NY). In addition, a titanium headplate (Luigs & Neuman, Ratingen, Germany) was fixed to the posterior base of the skull. A second layer of dental cement mixed with black carbon powder was used to reinforce and color the dental cement cap. Small pieces of electrical insulating tape were disposed like flower petals, glued, and cemented around the cement cap to limit light exposure coming from virtual reality screens during calcium imaging. Animals received a subcutaneous injection of meloxicam (2.5mg/kg) and were given post-surgical care for two days.

Animals were allowed to recover for 2-4 weeks with ad libitum water before beginning experiments. After at least one week of recovery, animals were anesthetized as described above and placed head-fixed under the 2-photon microscope (details below) to confirm presence of calcium signal and to determine a target area to image. The position (XYZ coordinates) of the chosen target area was noted relative to the cranial widow and surface of the brain.

### Virtual reality system and two-photon microscope

The virtual reality system (Luigs & Neumann, Ratingen, Germany) consisted of an arrangement of five monitors surrounding the sides and front of a treadmill belt (Fig. 1). Movement information of the belt was transmitted to a PC computer, which updated the position of the avatar in the virtual environment. A head fixation system and post were positioned at ∼45° angle behind the animal to avoid interfering with the display of the virtual reality environment and the microscope objective. A reward system (modular arm coupled with tubing and pump) was located at the side of the treadmill belt and adjusted for each animal. Virtual environment and behavioral tasks were custom-defined by modifying the pre-existing open-source code (Python 2.0) of the virtual reality system software (LN-treadmill-V4).

A 2-photon microscope (LSM 7MP, Carl Zeiss Ltd, Toronto, Canada) was installed above and posterior to the animal and was equipped with a water immersion long-working distance objective (20x; NA 1.0; WD 1.8mm). For EYFP and GCaMP6f epifluorescence imaging, a multi-LED light source (Colibri; Carl Zeiss) was used for illumination. For 2-photon calcium imaging, a Chameleon Ultra II laser (680-1080 nm wavelength; Coherent) was used with excitation set at 910 nm and emission filter set at 525-560nm bandpass.

The mouse position was adjusted under the microscope via micromanipulators (XY axes) of the virtual reality set-up that moved head-post, belt and reward systems with respect to the fixed microscope and monitors. Because the mouse was positioned on a treadmill belt, movement in the virtual environment was in forward or backward directions (see training section).

### Virtual Environment

The virtual reality environment for the spatial memory task consisted of a 360-cm long virtual corridor composed of three successive rooms, respectively starting at 67.6, 157.8 and 259.3 cm from the animal furthest start point (Fig. 1B). Rooms were, respectively, 22.5, 33.8 and 45.1 cm long, and 18.9, 28.0 and 32.5 cm wide. Walls of each room were covered with distinct symmetrical patterns (black triangle pattern on dark background, grey square grid on white background, and white dotted line on dark background). Rooms were separated by 9 cm wide corridors with the same asymmetrical pattern along the walls (left, grey honeycomb on dark background; right, black dot on grey background). Two target images were at each end of the corridor (start, dark squares on a white background; end, gray circles on white background). Several objects were present inside and outside the corridor with distinct patterns: brickwall (from the furthest start point X-position [X_pos_] = 78.9 cm, from avatar Y-position [Y_pos_] = 7.9 cm, corresponding to left of animal), ball (X_pos_ = 83.4 cm, Y_pos_ = -7.9 cm, corresponding to right of animal), tower 1 (outside, X_pos_ = 146.5 cm, Y_pos_ = -9 cm), cube (X_pos_ = 163.5 cm, Y_pos_ = -6.8 cm), cue card (X_pos_ = 180.4 cm, Y_pos_ = 13.8 cm), building (outside, X_pos_ = 246.9 cm, Y_pos_ = 15.8 cm), 3D square cross (X_pos_ = 268.3 cm, Y_pos_ = -6.8 cm), 3D diamond crystal (X_pos_ = 289.7 cm, Y_pos_ = 9.5 cm), pyramid (X_pos_ = 297.6 cm, Y_pos_ = -6.8 cm), moon (outside far away, X_pos_ = 2434.9 cm, Y_pos_ = -450.9 cm, Z-position [Z_pos_] = 450.9 cm), tower 2 (outside, X_pos_ = 338.2 cm, Y_pos_ = 11.3 cm). Two boxes, not visible from maze, were used to transiently teleport animals at the end of each trial. One box (X_pos_ = 180.4 cm, Y_pos_ = -225.5 cm) was completely black (ie. From the animal point of view, all screens black). The other box (X_pos_ = 180.4cm, Y_pos_ = -450.9 cm) was completely white (ie. From the animal point of view, all the screens white) and was used only after a failed trial (no reward).

### Habituation and training

Before behavioral experiments, mice were gently handled for 20 minutes for two days to habituate them to the experimenter and reduce stress related to experimental handling. After the post-operative recovery period of 2-4 weeks, mice were water restricted (1 ml/day) and their weight was controlled daily. At any stage, if animal weight decreased to less than 80% of the pre-restriction level, a larger amount of water was given (2-3 ml) for 1-2 days to restore weight. If animal weight remained under 80% of pre-restriction level, the animal was excluded, and ad libitum water access was restored.

During the first 2-3 days of water deprivation, mice were handled by the experimenter and habituated to equipment. Mice were gradually trained to run in the virtual reality setup. Initially, they were allowed to explore freely on the treadmill belt. Then, they were head-fixed and positioned on the belt in a black virtual environment for one habituation session, followed by 4 habituation sessions (2 of 5 min and 2 of 10 min duration) with reward (10% sweetened milk) available randomly in the environment (3-12 μl; separated by ≥5 sec) when animals licked the reward dispenser. The rationale was to habituate the animal to lick the reward dispenser while exploring the environment.

For the spatial learning task, animals were given 2 training sessions of 15 min per day in the virtual reality environment. Behavioral learning involved navigating the maze and learning the location of a reward zone at a specific location. Each trial consisted of navigating through the environment, stopping at the reward area (third room, X_pos_ 263.3 cm, length 28 cm), licking the dispenser for reward (available only once per trial), and then continuing to the end of the corridor to be teleported to the start for another trial. Reward was only given if the animal licked in the reward area. During the first 8 training sessions, reward was given directly after licking. After that, a reward delay was gradually introduced between the entrance in reward zone and licking. The delay was 0.5 sec for training sessions 9-10, then 1 sec for sessions 11-12, and finally 1.5 sec for sessions 13-16. In addition, to encourage animals to be precise, the amount of reward varied in the reward area; it was maximal at the center and decreased gradually in the surround (max 12 μl – min 3 μl).

When the animal arrived at the end of maze, there was two options for teleportation. After a success trial (reward obtained), the animal was teleported to the start of the maze by passing through a dark environment (black screens) for 2 seconds. After a failed trial (no reward obtained), the animal was teleported to the start by first passing through a well-lit environment (white screens) for 10 seconds followed by the black environment for 2 seconds. After 5 successive failed trials, the minimum reward was automatically given when the animal entered the reward zone. In addition, rewards were also given manually when animals performed too slowly for five trials (mostly in the first training sessions). Trials with given rewards were excluded from analysis. To encourage the use of visuals landmarks in the spatial learning task and not proprioceptive strategies, animals started each new trial at a random offset position (0-45 cm) from the start of the maze.

For the relearning task, the virtual environment was the same but the reward zone was relocated to another area of the maze located in the second room (X_pos_ 155.6 cm, length 28 cm). For the first 2 animals tested, the new reward zone was near the end of the first room (X_pos_ 57.7 cm, length 28 cm). In addition, for these animals the relearning task consisted of ten sessions (session 17-27) in the maze with a reward delay of 1.5 sec after entering the new reward zone. This relearning task and delay were judged too difficult and changed for all other animals to a relearning task organized similarly to the first learning task (no delay for sessions 17-24, 0.5 sec delay for sessions 25-26, 1 sec delay for sessions 27-28, and 1.5 sec delay for sessions 29-32).

### Recording procedures

Animals were handled for at least 10 min, placed on the belt and fixed to the headpost while screens were black. The imaging cannula cover was removed, and a small amount of distilled water was placed in it. The water immersion objective was lowered over the cranial window using epifluorescence imaging. The target area for imaging was identified relative to canula edge and brain surface using coordinates determined previously (see end of surgery section). Two-photon laser power illumination was set as in previous sessions and adjusted if necessary. Position of the reward dispenser was adjusted in front of the mouse to deliver reward. The virtual environment was then initialized, and calcium imaging and behavior were recorded for 15 min. After each session, water was removed from the canula with a small tissue and a new cover was placed above the canula.

### Data acquisition

Signals for time, belt movement sensor, lick sensor and pump status were digitized (30 Hz frequency) by the computer acquisition board of the virtual reality system and integrated with animal location in the virtual environment. Two-photon images were acquired with digital zoom (x2), at 5 Hz frequency, with 256 x 256-pixel resolution, throughout the 15 min training session. A 0.1sec TTL signal at the start of each frame acquisition was sent from the microscope output board to the virtual reality acquisition board and stored in the same csv file as the behavior variables (see below).

### Behavioral data analysis

Data analyses were performed by custom developed programs written in MATLAB (MathWorks). Data were directly acquired through the virtual reality system with a frequency of 30 Hz. Behavioral data were first re-interpolated at 100 Hz to have a fixed frequency.

#### Trials/min

A full trial corresponded to an animal that started at a random position, explored the total length of the corridor and reached the teleportation zone at end of the track. Number of trials per minute is computed from the total number of trials divided by total time of training session (15min).

#### Success ratio

The success ratio is the number of trials that the animal received a reward divided by the total number of trials multiplied by 100.

#### Lick in reward

Lick were considered only during trials. The percent of lick in reward zone corresponds to the number of licks in reward zone divided by the total number of licks multiplied by 100. If licking was absent, the value was set to zero.

#### Accuracy

The reward zone is subdivided is several sub zones (center of reward zone is more rewarded than the periphery (range 0.4 to 1.6 sec of pump activation). Accuracy of lick corresponds (in percent) to the sum of scores given as a function of reward duration (reflecting position in reward zone; no reward → 0 score; 0.4 sec reward → 0.25 score; 0.8 sec reward → 0.5 score; 1.2 sec reward → 0.75 score; 1.6 sec reward → 1 score) divided by the number of trials. If all rewards were in the center of the reward zone the mean accuracy was 100%, and if all trials were failures the accuracy was 0%.

#### Speed

Speed is computed from the distance traveled by the animal between two time points divided by the duration of the time interval. Speed measures were smoothed with an halfwidth of 100 points (1sec). For behavior analysis, mean speed during movement (speed > 1 cm x s^-1^) of the animal was calculated for each session.

#### Learning index

To quantitatively determine if animals learned the task, a composite learning index was calculated from the behavioral measures for each session. Learning index corresponds to the mean of the scores (between 0 and 1, with intervals of 0.25) for the 5 variables: i) success ratio in session ([0-20%] → 0 score; [20-40%] → 0.25 score; [40-60%] → 0.5 score; [60-80%] → 0.75 score; [80-100%] → 1 score); ii) running speed (cm/s; [0-5] → 0 score; [5-10] → 0.25 score; [10-15] → 0.5 score; [15-20] → 0.75 score; ≥20 ◊ 1 score); iii) trial duration (sec; >80 → 0 score; [60-80] → 0.25 score; [40-60] → 0.5 score; [20-40] → 0.75 score; [0-20] → 1 score); iv) accuracy in reward zone ([0-20%] → 0 score; [20-40%] → 0.25 score; [40-60%] → 0.5 score; [60-80%] → 0.75 score; [80-100%] → 1 score); and v) lick in reward zone ([0-20%] → 0 score; [20-40%] → 0.25 score; [40-60%] → 0.5 score; [60-80%] → 0.75 score; [80-100%] → 1 score).

### Calcium image processing

The time series analysis of 2-photon images consisted first of stabilizing images and then measuring changes in calcium fluorescence signals in specific regions of interest (ROI). Calcium signals of ROIs were then synchronized with behavior and re-interpolated at 100 Hz frequency for comparisons with behavioral measures.

#### Image stabilization

Raw movies with .czi extension were first converted to a .mat file containing raw movie and parameters of acquisition files, using bioformats and external code found at: bfopen:https://downloads.openmicroscopy.org/bio-formats/6.3.1/artifacts/bfmatlab.zip bfczifinfo: https://www.mathworks.com/matlabcentral/fileexchange/58666-read-information-from-zeiss-czi-image-file

Stabilization algorithm was strongly inspired by suite2P algorithm (66).

#### Reference frame

Stabilization consists of negating the drift in X-Y planes of an image from a reference frame. A temporary matrix made of 300-500 random frames in raw movie were computed. Then we computed a matrix correlation between all frames pair by pair. The reference frame corresponds to the mean frame of the first 20-30 best correlated frames.

#### Video stabilization

Frame drift from the reference frame was calculated using the phase correlation method involving fast Fourier transforms. Frame to be stabilized and reference frame were first transformed with fast Fourier transform to signals in frequency domain. The resulting complex number were then conjugated to give R number. The R number were then divided by the absolute value of itself and possible NaN values were replaced by 1. Finally, result components were transformed with inverse fast Fourier transform to find phase shift after reorganization of quarter. A maximum X-Y correction was set at 12% of image resolution (12*256/100 = 31 pixels). *Determination of region of interest (ROI).* To follow the same ROIs over successive sessions of training, a custom graphical user interface was coded to manually set and attribute ROIs. A square area including an individual SOM-IN was first determined, then using a manual threshold the main cell form was extracted and saved. A neuropil area corresponding to the 20 µm area around each ROI (without any other cell) was also determined.

#### Raw ROI, neuropil and corrected ROI fluorescence measures

For each frame, raw fluorescence of each ROI and neuropil was computed as the mean of fluorescence in the ROI cell mask region and associated neuropil mask region. Then ROI cell fluorescence was corrected by subtracting neuropil fluorescence with the constant k = 0,7.

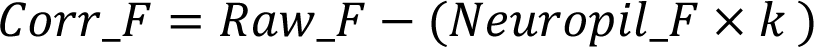

#### ΔF/F measures

For each ROI, fluorescence was calculated for each one-minute time window per session, and basal fluorescence (F0) was taken as the mean of fluorescence values under the 25^th^ percentile of Corr_F. Changes in fluorescence (ΔF/F) of ROIs were then calculated as:

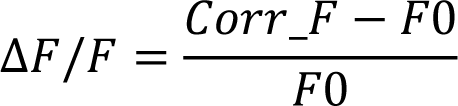

#### Out of ROI movement correction

After stabilization of X-Y movements, image movements in Z axis can occur. In the fluorescence signal, Z axis movement results in marked positive or negative changes in several ROIs with rapid kinetics that are different from normal GCaMP6f calcium signals. Z axis movements were detected using a positive and negative threshold equal to 3 times the standard deviation of the first derivative of ΔF/F. ΔF/F were corrected by interpolating values between points before and after the detected event.

#### Calcium signal synchronization

The TTL signal sent with every frame by the microscope to the virtual reality system acquisition card could be used to synchronize imaging and behavioral data. However, this required to launch imaging after initialization of behavior which was not systematically the case. To avoid problems, synchronization between imaging signals and behavior were aligned by checking several possible time drifts. A first-time drift was determined as the time of the first TTL signal present in behavior data. Second, if any teleportation in white box was present during a session, another putative time drift was computed using the cross correlation of global fluorescence (likely reflecting illumination of the white screens). Third, because mouse movement implies more instability of calcium images, another possible drift was calculated using the cross-correlation between speed vector and video X-Y correction vector. We considered that launching imaging and behavior recording was done under 10 sec. We first checked if time drifts 1 and 2 were similar (< 2 seconds). If it was the case, the TTL pulse was used for resynchronizing imaging and behaviors. However, if we found a difference between time drifts 1 and 2, and if the putative time drift 3 was inferior to 10 sec from the beginning of behavioral recording, then resynchronization was done using the time drift 3. Finally, if the time drifts 1 and 3 were both superior to 10 sec from beginning of behavioral recording, we used time drift 2 as beginning of recording. Resynchronization were then visually verified to confirm correct resynchronization.

### Analysis of calcium signal in relation to behavior

Several parameters were extracted from the analysis of the calcium signals for each ROI in relation to behavior during trials.

#### Speed correlation

Speed correlation was calculated by binning the speed measures in intervals of 5 cm*sec^-1^. For each ROI and trial, a correlation value was calculated between ΔF/F and the binned speed measure. In cases with less that 2 bins, speed correlation was not determined, and these trials and ROIs were excluded from the statistical analysis. A mean correlation value per ROI was obtained by averaging the correlation values for the trials. A p-value was obtained by comparing the distribution of correlation values for the trials against 0.

#### Acceleration and deceleration correlations

The acceleration and deceleration correlations were calculated using the first derivative of the speed measure and binning in intervals of 0.5m*s^2^. The acceleration (positive value) and deceleration (negative value) measures were considered separately because they may involve different mechanisms. For each ROI and trial, a correlation value was calculated between ΔF/F and the binned acceleration or deceleration measures. In cases with less that 2 bins, correlation was not determined, and these trials and ROIs were excluded from the statistical analysis. A mean correlation value for a ROI was obtained by averaging the correlation values for the trials. The p-value was obtained by comparing the distribution of correlation values for the trials against 0.

#### Place map

Place maps were calculated by binning the position signal into 100 bins (3.6cm per bin) per trial. In each bin, the average ΔF/F was calculated. The ΔF/F measures for each ROI were then organized by trials for each session. From this matrix, place map parameters were calculated: the normalized place map corresponds to the matrix with ΔF/F measures normalized between 0 (minimum) and 1 (maximum) for each trial. The mean place map corresponds to the matrix of ΔF/F measures averaged across all trials of a session. The mean normalized place map corresponds to the matrix of the normalized ΔF/F measures averaged across all trials of a session. *Place correlation.* The place correlation was calculated for each ROI by computing the correlation of ΔF/F measures between all trial pairs of the normalized place map in a session. The place correlation is the mean of the r distribution of all pairs in a session. To test for significance, we shuffled the position of the bins and calculated the mean r correlation for all pairs 1000 times. If the measured mean r was greater than the 95% confidence interval of the shuffled r distribution, the correlation was considered significant.

#### Activity correlation of ROIs

Activity correlation of ROIs was calculated per animal by computing the correlation of ΔF/F measures between all pairs of ROIs. The activity correlation of ROIs per animal is the mean of the r distribution of all pairs.

#### Modulation of ROI activity related to reward

The reward modulation of ROI activity was categorized as follows: For each ROI, using the mean normalized place map, ΔF/F measures were averaged for “before reward”, “in reward” and “after reward” locations. If the activity at “after reward” location was superior or inferior to 30% of the mean signal at “before reward” location (baseline), activity in the ROI was defined respectively as ‘sustained on’ and ‘sustained off’. If the activity at “in reward” location was superior or inferior to 30% of baseline, activity in the ROI was defined respectively as ‘transitory on’ and ‘transitory off’. Otherwise, activity in the ROI was considered non modulated.

#### Relearning

Animals did not get the same number of sessions during the relearning phase of training (10 sessions for the first 2 animals trained and 16 sessions for the subsequent 9 animals). To pool all data analysis, we compared parameters at 3 time points of relearning corresponding to the first session (start), the mid-point session (middle) and the last session (end) of relearning.

### Statistical analysis

All statistical analyses were conducted using Matlab codes (MathWorks). Before statistical tests, a Lilliefors goodness-of-fit test was used to verify data normality and a Levene test was used to test for equal variance. For multiple comparisons of repeated measures over time in the same animals, one-way ANOVA for repeated measures were used. If data normality and equal variance tests failed, Friedman tests were used. For data analysis with missing values, data rows involved were removed and the multi-comparison test was used if Friedman test was significant. For single pairwise comparison (ex. session 1 vs session 16) the p value was not corrected, but for multiple pairwise comparisons p values were corrected with Tukey-Kramer test.

For comparisons of two distributions, all available values were included, and Student t-tests were used. If normality or equal variance tests failed, the Wilcoxon rank sum test was used. For paired tests, all available values were included, and Student paired t-test were used. If normality test failed, the Wilcoxon signed rank test was used.

For comparison of multiple distributions, all available values were included, and a one-way ANOVA was used. If normality or equal variance test failed, the Kruskal Wallis test was used.

Results are expressed as mean ± s.e.m. in the figures and text. Details of all statistical tests are listed in Additional file 5: Table S1.

## Supplementary Information

**Additional file 1: Figure S1.** Inhibition of mTORC1 signaling in SOM-INs in SOM-Rptor-KO mice and other behavioral measures during learning. **A** Representative confocal immunofluorescence labelling of S6 phosphorylation (p-S6) in EYFP-expressing SOM-INs. Repeated mGluR1 stimulation (mGluR1/5 agonist DHPG in the presence of the mGluR5 antagonist MPEP) increased p-S6 in SOM-INs relative to sham-treatment in slices from control SOM-IRES-Cre mice (top) but not in slices from SOM-Rptor-KO mice (bottom). Scale bar: 20 µm. **B** Quantification of p-S6 immunofluorescence showing reduced basal level of p-S6 (sham-treatment) in SOM-INs of SOM-Rptor-KO mice (n=6; R.KO mice) relative to control SOM-IRES-Cre mice (n=6; Ctrl mice) suggesting deficit of constitutive mTORC1 activity. **C** Quantification of evoked p-S6 showing increased p-S6 in SOM-INs after repeated mGluR1 stimulation (DHPG in MPEP) relative to sham treatment of SOM-IRES-Cre mice (n=3 mice; Ctrl) but not in SOM-Rptor-KO mice (n=3 mice; R.KO) confirming a deficit in mTORC1 signaling in SOM-INs of SOM-Rptor-KO mice. **D, E** Summary plots of changes over training sessions in SOM-IRES-Cre (n=11 mice; Ctrl; black) and SOM-Rptor-KO (n=10 mice; R.KO; magenta) mice showing similar reduction in trial duration over training in both mice (**D**), and increase in percentage time spent in reward zone over training only in control SOM-IRES-Cre mice (**E**), indicative of a spatial learning deficit in SOM-Rptor-KO mice. Details of statistical tests provided in Additional file 5: Table S1. * p<0.05, *** p<0.001, ns not significant.

**Additional file 2: Figure S2.** Deceleration correlation over training, and speed, acceleration, and deceleration correlations with mean learning index. **A** Mean correlation of Ca^2+^ activity with deceleration for all SOM-INs decreased at the end relative to the start of training in control but not in SOM-Rptor-KO mice. **B** Mean speed correlation as a function of mean learning index for all animals, showing correlation in control but not SOM-Rptor-KO mice. **C** Mean acceleration correlation as a function of mean learning index for all animals, showing absence of correlation. **D** Mean deceleration correlation as a function of mean learning index for all animals, showing correlation in control but not SOM-Rptor-KO mice. **E** Mean speed correlation as a function of mean place correlation for all animals, showing correlation in control but not SOM-Rptor-KO mice. Details of statistical tests provided in Additional file 5: Table S1. * p<0.05, ** p<0.01, *** p<0.001, ns not significant.

**Additional file 3: Figure S3.** SOM-INs with no response modulation; deceleration correlation of response types; and activity correlation between cells. **A** Three representative examples of SOM-IN responses with no modulation related to reward location. Top: mean Ca^2+^ responses (grey) and speed (red) as function of position for all trials in a session with reward zone indicated in green. Bottom: color-coded Ca^2+^ activity in each trial of the session. **B** Similar representation as in Fig. 3F of Ca^2+^ activity correlation with deceleration for SOM-INs with different response types, showing response-specific changes over training. **C** Mean activity correlation between all SOM-INs, showing less correlation in control mice relative to SOM-Rptor-KO mice. Details of statistical tests provided in Additional file 5: Table S1. * p<0.05, ** p<0.01, *** p<0.001, ns not significant.

**Additional file 4: Figure S4.** Examples of remapping of SOM-IN activity, activity correlation, and response type distribution during relearning in control and SOM-Rptor-KO mice. **A** Example of no remapping of SOM-IN responses for a cell with “reward off transient” responses at both end of learning (left) and end of relearning (right). For each session, top left is mean Ca^2+^ responses (grey) and speed (red) as function of position for all trials in the session with reward zone indicated (green for learning; red for relearning); bottom left is color-coded Ca^2+^ activity in each trial of the session; top right is place correlation matrix of all paired laps; and bottom right is distribution of r values (blue), mean r (red) versus r distribution obtained by shuffling position measures (gray). **B** Similar representation of remapping for a SOM-IN with “reward on sustained” response at end of learning and “non-modulated” response at end of relearning. **C** Similar representation of remapping for a SOM-IN with “reward off transient” response at end of learning and “reward off sustained” response at end of relearning. **D** Similar representation of remapping for a SOM-IN with a “non-modulated” response at end of learning and “reward on transient” response at end of relearning. **E** Mean speed correlation with activity for all SOM-INs showing no change during relearning in control (n=40 cells) and SOM-Rptor-KO (n=32 cells) mice. **F** Mean acceleration correlation with activity for all SOM-INs showing no change during relearning in control and SOM-Rptor-KO mice. **G** Mean deceleration correlation with activity for all SOM-INs showing a decrease during relearning in control but not SOM-Rptor-KO mice. **H** Cell response identity matrix for all cells in control mice (grey top) and SOM-Rptor-KO mice (magenta, bottom) during learning (green, left) and relearning (red, right) ordered by response type at end of learning, showing a gradual acquisition of a new spatial coding related to reward relocation during relearning in control but not SOM-Rptor-KO mice. **I** Distribution of cells with different response types at start, middle and end of learning and relearning for control mice (grey, left) and SOM-Rptor-KO mice (magenta, right), showing a decrease in number of modulated cells at start of relearning relative to end of learning, and an increase during relearning in control but not SOM-Rptor-KO mice. Details of statistical tests provided in Additional file 5: Table S1. * p<0.05, ns not significant.

**Additional file 5: Table S1.** Details of statistical tests.

## Abbreviations

ACSF: artificial cerebrospinal fluid
DHPG: (S)-3,5-dihydroxyphenylglycine
EYFP: enhanced yellow fluorescent protein
IN: inhibitory interneuron
LTP: long-term potentiation
mGluR1a: metabotropic glutamate receptor 1a
MPEP: 2-methyl-6 (phenylethynyl)-pyridine
mTORC1: mechanistic target of rapamycin complex 1
NS-SOM: narrow spike SOM neurons
O-LM: oriens-lacunosum/moleculare
PBS: phosphate-buffered saline
PC: CA1 pyramidal cell
Raptor: regulatory-associated protein of mTOR
ROI: region of interest
SOM-IN: CA1 Somatostatin interneuron
SOM-IRES-Cre: *Sst*^ires-Cre/wt^
SOM-Rptor-KO: *Sst*^ires-Cre/wt^;*Rptor*^fl/fl^
TORO: theta-OFF/ripple ON
TSC1: tuberous sclerosis complex 1
VIP: vasoactive intestinal peptide
WS-SOM: wide spike SOM neuron

## Acknowledgments

The authors thank Nicolas Stifani and Joanne Vallée for assistance with 2-photon microscope maintenance.

## Author contributions

François-Xavier Michon: Conceptualization, Methodology, Software, Validation, Formal analysis, Investigation, Data curation, Writing – original draft, Writing – review and editing, Visualization. Isabel Laplante: Conceptualization, Methodology, Validation, Formal analysis, Investigation, Data curation, Writing – original draft, Writing – review and editing, Visualization. Anthony Bosson: Methodology, Writing – review and editing. Richard Robitaille: Conceptualization, Methodology, Resources, Writing – review and editing. Jean-Claude Lacaille: Conceptualization, Methodology, Validation, Formal analysis, Resources, Writing – original draft, Writing – review and editing, Visualization, Supervision, Project administration, Funding acquisition.

## Funding

This work was supported by a grant to J.C.L. from the Canadian Institutes of Health Research (CIHR; PJT-153311), grants to R.R. from CIHR (PJT-173415) and from the Natural Sciences and Engineering Research Council of Canada (RGPIN-2020-05264), by infrastructure grants to J.C.L. and R.R. from the Canada Foundation for Innovation, and by a Research Centre grant to J.C.L. and R.R. (Centre Interdisciplinaire de Recherche sur le Cerveau et l’Apprentissage; CIRCA) from the Fonds de la Recherche du Québec – Santé (FRQS). J.C.L. is the recipient of the Canada Research Chair in Cellular and Molecular Neurophysiology (CRC 950-231066). J.C.L and R.R. are members of the Research Group on Neural Signaling and Circuitry (GRSNC) at Université de Montréal.

## Availability of data and materials

The data and analysis for the work presented in the current study are available from the corresponding author upon reasonable request.

## Declarations

### Consent for publication

All authors have given their consent for publication.

### Competing interests

The authors declare that they have no competing interests.

## Notes

### Competing Interest Statement

The authors have declared no competing interest.

