## Supplementary figures and images for "mTORC1-mediated acquisition of reward-related spatial representations by hippocampal somatostatin interneuronsa"

### Supplemental Fig S1

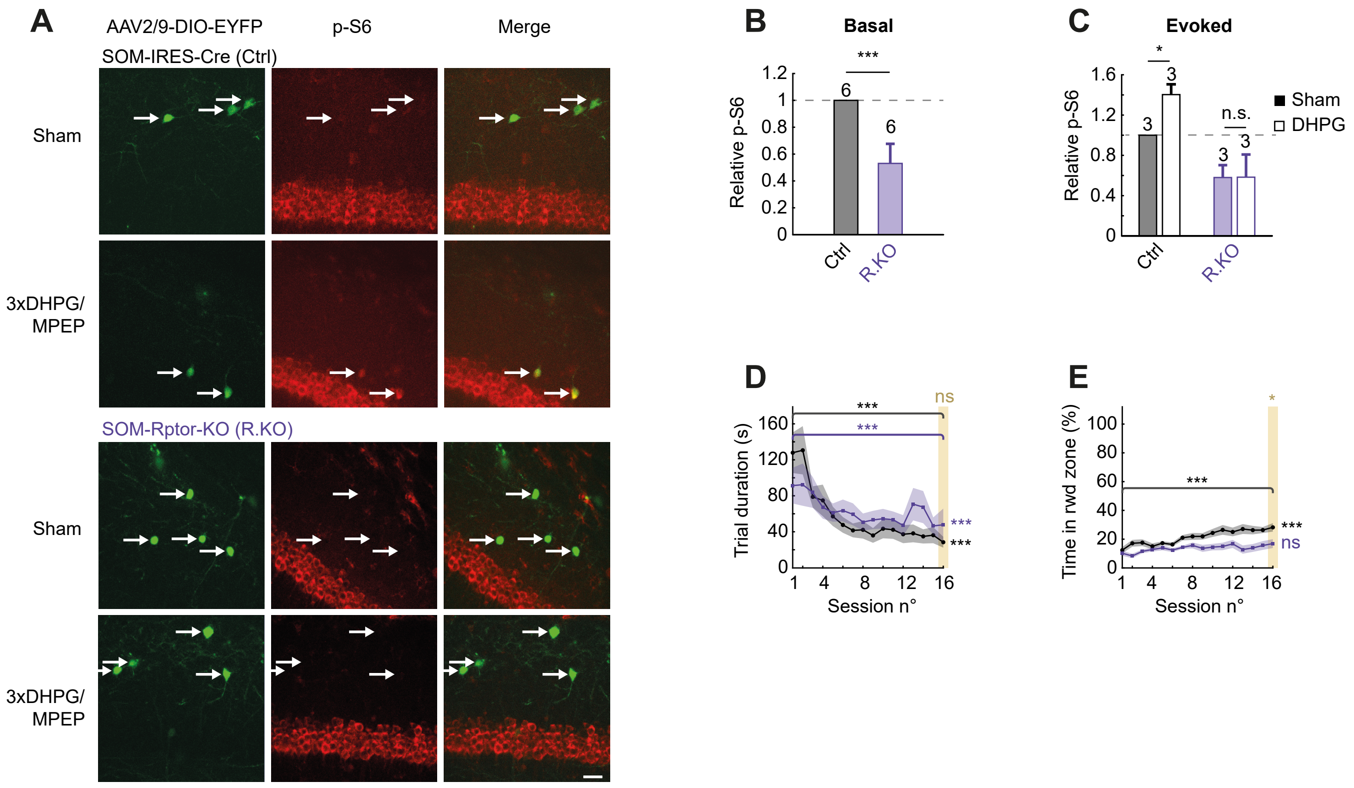

### Supplemental Fig S2

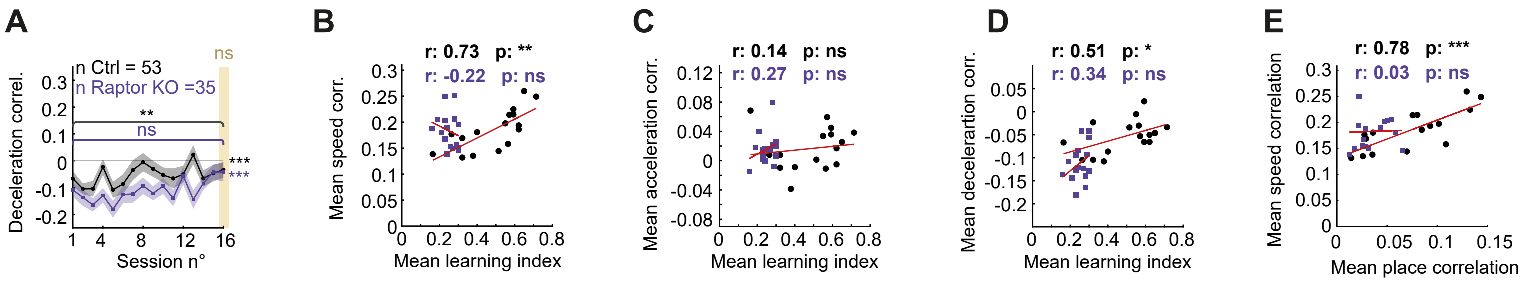

### Supplemental Fig S3

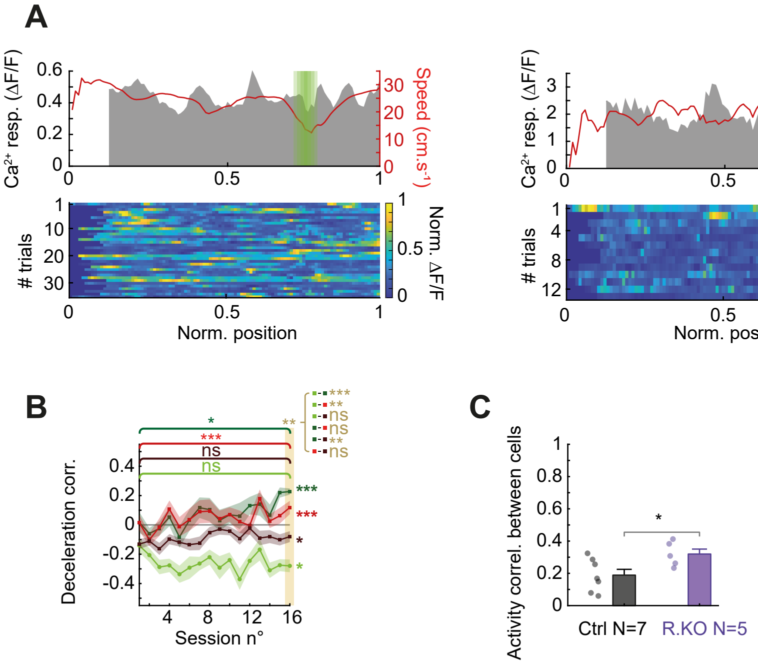

### Supplemental Fig S4

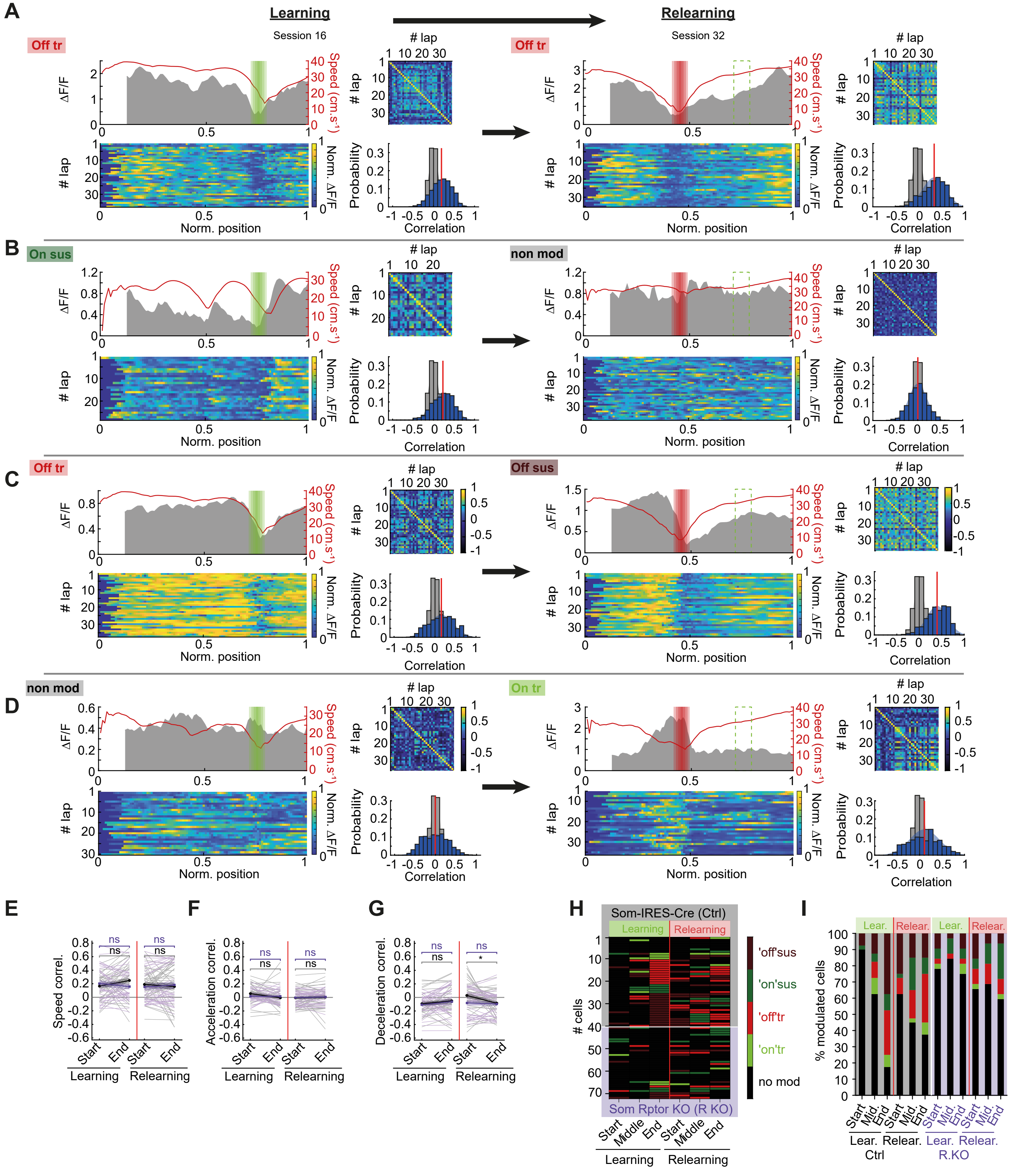
