## Supplemental Table S1 for "mTORC1-mediated acquisition of reward-related spatial representations by hippocampal somatostatin interneuronsa"

**Table S1 - Details of statistical tests**

| Figure | Panel | Data name | Test | Group size | Statistic | P value | Multicomparison test | Statistic | Pvalues2 |
|---|---|---|---|---|---|---|---|---|---|
| 1 | F | Trials/min | Friedman's test | Som-Ctrl n=11 | Sum of Square(df:15) =1905,31818181818 Chi-sq = 84,1143812709030 | 1,22586E-11 | only S1-S16 comparison: no correction | T= -10,9545454545455 | 6,7415E-08 |
| 1 | F | Trials/min | Friedman's test | Som-Rptor KO n=10 | Sum of Square(df:15) =1174,400 Chi-sq = 51,8880706921944 | 5,8943E-06 | only S1-S16 comparison: no correction | T= -8,15000000000000 | 0,000127824 |
| 1 | F | Trials/min | unpaired t-test | S1 :Som-Ctrl n=11 vs Som-Rptor KO n=10 | t(df:19)=-1,43553000482949 | 0,167395217 | | | |
| 1 | F | Trials/min | unpaired t-test | s16: Som-Ctrl n=11 vs Som-Rptor KO n=10 | t(df:19)=1,68512154890916 | 0,108319879 | | | |
| 1 | G | Speed | Friedman's test | Som-Ctrl n=11 | Sum of Square(df:15) = 2344,18181818182 Chi-sq = 103,461281262538 | 2,87011E-15 | only S1-S16 comparison: no correction | T=-11,7272727272727 | 7,5629E-09 |
| 1 | G | Speed | Friedman's test | Som-Rptor KO n=10 | Sum of Square(df:15) = 1331,15000000000 Chi-sq = 58,7358435063980 | 4,15437E-07 | only S1-S16 comparison: no correction | T = -9,70000000000000 | 5,2108E-06 |
| 1 | G | Speed | unpaired t-test | S1 :Som-Ctrl n=11 vs Som-Rptor KO n=10 | t(df:19)= -1,35233044732337 | 0,192147329 | | | |
| 1 | G | Speed | unpaired t-test | s16: Som-Ctrl n=11 vs Som-Rptor KO n=10 | t(df:19)= 1,23601872741081 | 0,23151513 | | | |
| 1 | H | % Success trials | Friedman's test | Som-Ctrl n=11 | Sum of Square(df:15) = 733,590909090909 Chi-sq = 32,4771934531795 | 0,005540512 | only S1-S16 comparison: no correction | T = -6,81818181818182 | 0,000767004 |
| 1 | H | % Success trials | Friedman's test | Som-Rptor KO n=10 | Sum of Square(df:15) = 461,100000000000 Chi-sq = 22,5734334203655 | 0,093626198 | | | |
| 1 | H | % Success trials | unpaired t-test | S1 :Som-Ctrl n=11 vs Som-Rptor KO n=10 | t(df:19)= 1,64069215277192 | 0,117311856 | | | |
| 1 | H | % Success trials | Wilcoxon Rank test | s16: Som-Ctrl n=11 vs Som-Rptor KO n=10 | z-val(ranksum:141)= 1,38215299079 | 0,166924731 | | | |
| 1 | I | % Lick in rwd zone | Friedman's test | Som-Ctrl n=11 | Sum of Square(df:15) = 1855,22727272727 Chi-sq = 81,8592057761733 | 3,18778E-11 | only S1-S16 comparison: no correction | T=-11 | 5,99748E-08 |
| 1 | I | % Lick in rwd zone | Friedman's test | Som-Rptor KO n=10 | Sum of Square(df:15) = 340,350000000000 Chi-sq = 16,1814580031696 | 0,370095448 | | | |
| 1 | I | % Lick in rwd zone | Wilcoxon Rank test | S1 :Som-Ctrl n=11 vs Som-Rptor KO n=10 | z-val(ranksum:131)= 0,67670386314 | 0,498593865 | | | |
| 1 | I | % Lick in rwd zone | Wilcoxon Rank test | s16: Som-Ctrl n=11 vs Som-Rptor KO n=10 | z-val(ranksum:145)= 1,65535769809 | 0,037522733 | | | |
| 1 | J | % Accuracy | Friedman's test | Som-Ctrl n=11 | Sum of Square(df:15) = 802,136363636364 Chi-sq = 35,4547281007233 | 0,002117492 | only S1-S16 comparison: no correction | T= -6,63636363636364 | 0,001067577 |
| 1 | J | % Accuracy | Friedman's test | Som-Rptor KO n=10 | Sum of Square(df:15) = 264,700000000000 Chi-sq = 12,9100959193627 | 0,609242051 | | | |
| 1 | J | % Accuracy | unpaired t-test | S1 :Som-Ctrl n=11 vs Som-Rptor KO n=10 | t(df:19)= 1,49395356123137 | 0,151609979 | | | |
| 1 | J | % Accuracy | Wilcoxon Rank test | s16: Som-Ctrl n=11 vs Som-Rptor KO n=10 | z-val(ranksum:142)= 1,45303263134 | 0,146214694 | | | |
| 1 | K | Learning index | Friedman's test | Som-Ctrl n=11 | Sum of Square(df:15) = 2143,22727272727 Chi-sq = 98,2039711191335 | 2,85419E-14 | only S1-S16 comparison: no correction | T = -12,5909090909091 | 2,60291E-10 |
| 1 | K | Learning index | Reapeated ANOVA 1w | Som-Rptor KO n=10 | Sum of Square(df:15) = 1,02793750000000 F = 4,54364908834183 | 6,20836E-07 | only S1-S16 comparison: no correction | T = -0,290000000000000 | 0,003758319 |
| 1 | K | Learning index | unpaired t-test | S1 :Som-Ctrl n=11 vs Som-Rptor KO n=10 | t(df:19)= 0,0381576419620768 | 0,969960071 | | | |
| 1 | K | Learning index | unpaired t-test | s16: Som-Ctrl n=11 vs Som-Rptor KO n=10 | t(df:19)= 2,33339029384077 | 0,030767771 | | | |
| Sup1 | B | Relative p-s6 | unpaired t-test | Som-Ctrl n=6 vs Som-Rptor KO n=6 | t(df:5) = 7,7953 | 0,00055648 | | | |
| Sup1 | C | Relative p-s6 | unpaired t-test | Som-Ctrl Basal n=3 vs Som-Ctrl DHPG n=3 | t(df:2) = 6,8393 | 0,0197 | | | |
| Sup1 | C | Relative p-s6 | unpaired t-test | Som-Rptor KO Basal n=3 vs Som-Rptor KO DHPG n=3 | t(df:2) = 3,2072 | 0,0873 | | | |
| Sup1 | D | Trials duration | Friedman's test | Som-Ctrl n=11 | Sum of Square(df:15) = 1945,81818181818 Chi-sq = 85,8449197860963 | 5,87011E-12 | only S1-S16 comparison: no correction | T = 10,1818181818182 | 5,29001E-07 |
| Sup1 | D | Trials duration | Friedman's test | Som-Rptor KO n=10 | Sum of Square(df:15) = 1097,65000000000 Chi-sq = 48,4328577732019 | 2,16492E-05 | only S1-S16 comparison: no correction | T = 8,60000000000000 | 5,35781E-05 |
| Sup1 | D | Trials duration | unpaired t-test | S1 :Som-Ctrl n=11 vs Som-Rptor KO n=10 | t(df:19)= 1,14555966171812 | 0,266202021 | | | |
| Sup1 | D | Trials duration | Wilcoxon Rank test | s16: Som-Ctrl n=11 vs Som-Rptor KO n=10 | z-val(ranksum:111)= -0,66896985070 | 0,50351471 | | | |
| Sup1 | E | % Time in rwd zone | Reapeated ANOVA 1w | Som-Ctrl n=11 | Sum of Square(df:15) = 4091,91473705755 F = 8,00887762932794 | 4,48575E-13 | only S1-S16 comparison: no correction | T = -15,8210246006651 | 7,54E-06 |
| Sup1 | E | % Time in rwd zone | Friedman's test | Som-Rptor KO n=10 | Sum of Square(df:15) = 523,800000000000 Chi-sq =23,1088235294118 | 0,081856517 | | | |
| Sup1 | E | % Time in rwd zone | unpaired t-test | S1 :Som-Ctrl n=11 vs Som-Rptor KO n=10 | t(df:19)= 0,716775737362498 | 0,482231356 | | | |
| Sup1 | E | % Time in rwd zone | unpaired t-test | s16: Som-Ctrl n=11 vs Som-Rptor KO n=10 | t(df:19)= 2,51282711742283 | 0,021159953 | | | |
| 2 | G | Place correlation | Friedman's test | Som-Ctrl n=44 | Sum of Square(df:15) = 3057,04545454546 Chi-sq = 134,869652406417 | 2,35682E-21 | only S1-S16 comparison: no correction | T=-4,09090909090909 | 5,57058E-05 |
| 2 | G | Place correlation | Friedman's test | Som-Rptor KO n=32 | Sum of Square(df:15) = 692,750000000000 Chi-sq = 30,5625000000000 | 0,010047235 | only S1-S16 comparison: no correction | T = -1,90625000000000 | 0,109250653 |
| 2 | G | Place correlation | Wilcoxon Rank test | S1 :Som-Ctrl n=47 vs Som-Rptor KO n=32 | z-val(ranksum:1816)= -0,6341550229 | 0,525979657 | | | |
| 2 | G | Place correlation | Wilcoxon Rank test | s16: Som-Ctrl n=53 vs Som-Rptor KO n=35 | z-val(ranksum:2761)= 3,42728101940 | 0,000609658 | | | |
| 2 | H | Speed correlation | Friedman's test | Som-Ctrl n=37 | Sum of Square(df:15) = 685,729729729730 Chi-sq = 30,2527821939587 | 0,011041932 | only S1-S16 comparison: no correction | T=-1,72972972972973 | 0,118127685 |
| 2 | H | Speed correlation | Friedman's test | Som-Rptor KO n=25 | Sum of Square(df:15) = 651,120000000000 Chi-sq = 28,7258823529412 | 0,017442559 | only S1-S16 comparison: no correction | T = -0,200000000000000 | 0,881930721 |
| 2 | H | Speed correlation | Wilcoxon Rank test | S1 :Som-Ctrl n=39 vs Som-Rptor KO n=32 | z-val(ranksum:1320)= -0,96494721637 | 0,33457125 | | | |
| 2 | H | Speed correlation | Wilcoxon Rank test | s16: Som-Ctrl n=53 vs Som-Rptor KO n=35 | z-val(ranksum:2591)= 1,97793332460 | 0,047936229 | | | |
| 2 | I | Acceleration correlation | Friedman's test | Som-Ctrl n=37 | Sum of Square(df:15) = 910,432432432433 Chi-sq = 40,1661367249603 | 0,000428031 | only S1-S16 comparison: no correction | T = 2,05405405405405 | 0,063498686 |
| 2 | I | Acceleration correlation | Reapeated ANOVA 1w | Som-Rptor KO n=25 | Sum of Square(df:15) = 0,147570052078411 F = 1,71886708402397 | 0,045474817 | only S1-S16 comparison: no correction | T = -0,0133057849684411 | 0,638876124 |
| 2 | I | Acceleration correlation | Wilcoxon Rank test | S1 :Som-Ctrl n=39 vs Som-Rptor KO n=32 | z-val(ranksum:1574)= 1,95878506793 | 0,050137963 | | | |
| 2 | I | Acceleration correlation | Wilcoxon Rank test | s16: Som-Ctrl n=53 vs Som-Rptor KO n=35 | z-val(ranksum:2339)= -0,16198591888 | 0,871316943 | | | |
| 2 | J | place corr. vs learning in | linear correlation | Som-Ctrl n=16 | $r^2$=0,77 | 8,60232E-06 | | | |
| 2 | J | place corr. vs learning in | linear correlation | Som-Rptor KO n=16 | $r^2$=-0,01 | 0,756482626 | | | |
| Sup2 | A | Deceleration correlation | Friedman's test | Som-Ctrl n=37 | Sum of Square(df:15) = 1207,40540540541 Chi-sq = 53,2678855325914 | 3,47988E-06 | only S1-S16 comparison: no correction | T = -3,18918918918919 | 0,003961732 |
| Sup2 | A | Deceleration correlation | Friedman's test | Som-Rptor KO n=25 | Sum of Square(df:15) =1314,32000000000 Chi-sq = 57,9847058823530 | 5,58121E-07 | only S1-S16 comparison: no correction | T = -0,720000000000001 | 0,59287138 |
| Sup2 | A | Deceleration correlation | Wilcoxon Rank test | S1 :Som-Ctrl n=39 vs Som-Rptor KO n=32 | z-val(ranksum:1432)= 0,31779698741 | 0,750638936 | | | |
| Sup2 | A | Deceleration correlation | unpaired t-test | s16: Som-Ctrl n=53 vs Som-Rptor KO n=35 | t(df:86)= 0,189226304529002 | 0,850361323 | | | |
| Sup2 | B | speed corr vs learning in | linear correlation | Som-Ctrl n=16 | $r^2$=0,54 | 0,001181643 | | | |
| Sup2 | B | speed corr vs learning in | linear correlation | Som-Rptor KO n=16 | $r^2$=0,05 | 0,405828541 | | | |
| Sup2 | C | accel. corr vs learning in | linear correlation | Som-Ctrl n=16 | $r^2$=0,02 | 0,596293979 | | | |
| Sup2 | C | accel. corr vs learning in | linear correlation | Som-Rptor KO n=16 | $r^2$=0,07 | 0,307348255 | | | |
| Sup2 | D | decel. corr vs learning in | linear correlation | Som-Ctrl n=16 | $r^2$=0,26 | 0,044852146 | | | |
| Sup2 | D | decel. corr vs learning in | linear correlation | Som-Rptor KO n=16 | $r^2$=0,12 | 0,199593665 | | | |
| Sup2 | E | speed corr vs place corr | linear correlation | Som-Ctrl n=16 | $r^2$=0,61 | 0,000366788 | | | |
| Sup2 | E | speed corr vs place corr | linear correlation | Som-Rptor KO n=16 | $r^2$=0,00 | 0,907776076 | | | |
| 3 | E | % modulated cells | paired t-test | Som-Ctrl n=7 Vs Som-Ctrl n=7 | t(df:6) = -3,7587 | 0,0094134 | | | |
| 3 | E | % modulated cells | paired t-test | Som-Rptor KO n=5 Vs Som-Rptor KO n=5 | t(df:4) = -0,35863 | 0,738 | | | |
| 3 | E | % modulated cells | unpaired t-test | Som-Ctrl n=7 Vs Som-Rptor KO n=5 | t(df:10) = 2,4855 | 0,032239 | | | |
| 3 | F | Place corr. | Reapeated ANOVA 1w | "On transient" n=5 | Sum of Square(df:15) = 0,115497905064709 F = 2,75846262440902 | 0,002750118 | only S1-S16 comparison: no correction | T = -0,0720982246751117 | 0,171051844 |
| 3 | F | Place corr. | Friedman's test | "On sustained" n=6 | Sum of Square(df:15) = 828,333333333333 Chi-sq = 36,5441176470588 | 0,001474579 | only S1-S16 comparison: no correction | T = -9,83333333333333 | 0,000347028 |
| 3 | F | Place corr. | Friedman's test | "Off transient" n=15 | Sum of Square(df:15) = 1788,66666666667 Chi-sq = 78,9117647058824 | 1,10377E-10 | only S1-S16 comparison: no correction | T = -5,06666666666667 | 0,003562965 |

| | | | | | | | | |
|---|---|---|---|---|---|---|---|---|
| 3 | F | Place corr. | Friedman's test | "Off sustained" n=17 | Sum of Square(df:15) =<br>1375,76470588235<br>Chi-sq = 60,6955017301038 | 1,91434E-07 | only S1-S16 comparison: no correction | T = -3,52941176470588 | 0,030671055 |
| 3 | F | Place corr. | Kruskal-Wallis test | S1 : "On transient" n=5 vs<br>"On sustained" n=6 vs<br>"Off transient" n=15 vs<br>"Off sustained" n=19 | Sum of Square(df:3) =<br>850,477192982456<br>Chi-sq = 4,93030256801424 | 0,176972455 | | | |
| 3 | F | Place corr. | Kruskal-Wallis test | S16 : "On transient" n=6 vs<br>"On sustained" n=8 vs<br>"Off transient" n=15 vs<br>"Off sustained" n=19 | Sum of Square(df:3) =<br>46,5600877192982<br>Chi-sq = 0,237551467955603 | 0,971310962 | | | |
| 3 | G | Speed corr. | Reapeated ANOVA 1w | "On transient" n=5 | Sum of Square(df:15) =<br>0,733759804283694<br>F = 1,97740495774470 | 0,032445887 | only S1-S16 comparison: no correction | T = 0,282380826159446 | 0,045344412 |
| 3 | G | Speed corr. | Friedman's test | "On sustained" n=6 | Sum of Square(df:15) =<br>299,333333333333<br>Chi-sq = 13,2058823529412 | 0,586399228 | | | |
| 3 | G | Speed corr. | Friedman's test | "Off transient" n=12 | Sum of Square(df:15) = 1500<br>Chi-sq = 66,1764705882353 | 2,12343E-08 | only S1-S16 comparison: no correction | T = -6,33333333333333 | 0,001120135 |
| 3 | G | Speed corr. | Friedman's test | "Off sustained" n=17 | Sum of Square(df:15) =<br>479,294117647059<br>Chi-sq = 21,1453287197232 | 0,13224993 | | | |
| 3 | G | Speed corr. | ANOVA n-way | S1 : "On transient" n=5 vs<br>"On sustained" n=6 vs<br>"Off transient" n=12 vs<br>"Off sustained" n=19 | Sum of Square(df:3) =<br>0,196923925148205<br>F = 1,46773121301300 | 0,23867201 | | | |
| 3 | G | Speed corr. | ANOVA n-way | S16 : "On transient" n=6 vs<br>"On sustained" n=8 vs<br>"Off transient" n=15 vs<br>"Off sustained" n=19 | Sum of Square(df:3) =<br>1,49321457347455<br>F = 12,5725612873868 | 4,55733E-06 | multicomparison tukey-kramer correction | Constant' coeffs=0,152455832371639<br>''on''tr' coeffs=-0,216922808756565<br>'on''sus' coeffs= -0,173602493317361<br>''off''tr' coeffs= 0,197348101752208<br>''off''sus' coeffs= 0,193177200321718 | "On transient" vs "On sustained" p: 0,977563911977934<br>"On transient" vs "Off transient" p: 0,000508280994916685<br>"On transient" vs "Off sustained" p: 0,000381954187290178<br>"On sustained" vs "Off transient" p: 0,000597459945580504<br>"On sustained" vs "Off sustained" p: 0,000416553551710702<br>"Off transient" vs "Off sustained" p: 0,999918463895285 |
| 3 | H | Acceleration corr. | Reapeated ANOVA 1w | "On transient" n=5 | Sum of Square(df:15) =<br>0,158035121016093<br>F = 1,83254705069099 | 0,050594321 | | | |
| 3 | H | Acceleration corr. | Friedman's test | "On sustained" n=6 | Sum of Square(df:15) = 517<br>Chi-sq = 22,8088235294118 | 0,088284096 | | | |
| 3 | H | Acceleration corr. | Friedman's test | "Off transient" n=12 | Sum of Square(df:15) =<br>1247,33333333333<br>Chi-sq = 55,0294117647059 | 1,76565E-06 | only S1-S16 comparison: no correction | T = 4,16666666666667 | 0,032054341 |
| 3 | H | Acceleration corr. | Friedman's test | "Off sustained" n=17 | Sum of Square(df:15) =<br>559,529411764706<br>Chi-sq = 24,6851211072664 | 0,054335325 | | | |
| 3 | H | Acceleration corr. | ANOVA n-way | S1 : "On transient" n=5 vs<br>"On sustained" n=6 vs<br>"Off transient" n=12 vs<br>"Off sustained" n=19 | Sum of Square(df:3) =<br>0,212189861364568<br>F = 4,69418488808290 | 0,006947459 | multicomparison tukey-kramer correction | Constant' coeffs=0,0380571389031335<br>''on''tr' coeffs=-0,106107534196297<br>'on''sus' coeffs= -0,0408763519367231<br>''off''tr' coeffs=0,116358954179368<br>''off''sus' coeffs=0,0306249319536519 | "On transient" vs "On sustained" p: 0,816380518034974<br>"On transient" vs "Off transient" p: 0,00821646641392044<br>"On transient" vs "Off sustained" p: 0,137173492855312<br>"On sustained" vs "Off transient" p: 0,0663346795902567<br>"On sustained" vs "Off sustained" p: 0,603434505133967<br>"Off transient" vs "Off sustained" p: 0,247571356711219 |
| 3 | H | Acceleration corr. | Kruskal-Wallis test | S16 : "On transient" n=6 vs<br>"On sustained" n=8 vs<br>"Off transient" n=15 vs<br>"Off sustained" n=19 | Sum of Square(df:3) =<br>2289,90219298246<br>Chi-sq = 11,6831744539921 | 0,008551156 | multicomparison tukey-kramer correction | 'on''tr' mnRank(n:6)=12,8333333333333<br>'on''sus' mnRank(n:8)=37,3750000000000<br>''off''tr' mnRank(n:15)=25,6000000000000<br>''off''sus' mnRank(n:19)=21,8947368421053 | "On transient" vs "On sustained" p: 0,00643295421301089<br>"On transient" vs "Off transient" p: 0,233215277674377<br>"On transient" vs "Off sustained" p: 0,510570324383959<br>"On sustained" vs "Off transient" p: 0,219068691669257<br>"On sustained" vs "Off sustained" p: 0,0431897307930264<br>"Off transient" vs "Off sustained" p: 0,869663650761245 |
| Sup3 | B | Deceleration corr. | Reapeated ANOVA 1w | "On transient" n=5 | Sum of Square(df:15) =<br>0,308014257209970<br>F = 1,96539540849005 | 0,033673396 | only S1-S16 comparison: no correction | T = 0,151235252043244 | 0,087844195 |
| Sup3 | B | Deceleration corr. | Reapeated ANOVA 1w | "On sustained" n=6 | Sum of Square(df:15) =<br>0,779147444708772<br>F = 3,27650228974189 | 0,000333503 | only S1-S16 comparison: no correction | T = -0,215417582338703 | 0,030576328 |
| Sup3 | B | Deceleration corr. | Friedman's test | "Off transient" n=12 | Sum of Square(df:15) =<br>1023,00000000000<br>Chi-sq = 45,1323529411765 | 7,29756E-05 | only S1-S16 comparison: no correction | T = -6,75000000000000 | 0,00051497 |
| Sup3 | B | Deceleration corr. | Friedman's test | "Off sustained" n=17 | Sum of Square(df:15) =<br>593,882352941176<br>Chi-sq = 26,2006920415225 | 0,035965394 | only S1-S16 comparison: no correction | T = -2,82352941176471 | 0,083799863 |
| Sup3 | B | Deceleration corr. | ANOVA n-way | S1 : "On transient" n=5 vs<br>"On sustained" n=6 vs<br>"Off transient" n=12 vs<br>"Off sustained" n=19 | Sum of Square(df:3) =<br>0,227636786429816<br>F = 2,68227999947513 | 0,060393025 | | Constant' coeffs=-0,0602530111647485<br>''on''tr' coeffs=-0,0811266521749768<br>'on''sus' coeffs= 0,0768994689187275<br>''off''tr' coeffs=0,0750418040533553<br>''off''sus' coeffs=-0,0708146207971061 | |
| Sup3 | B | Deceleration corr. | Kruskal-Wallis test | S16 : "On transient" n=6 vs<br>"On sustained" n=8 vs<br>"Off transient" n=15 vs<br>"Off sustained" n=19 | Sum of Square(df:3) =<br>4838,81052631579<br>Chi-sq = 24,6878088077336 | 1,79441E-05 | multicomparison tukey-kramer correction | on''tr' mnRank(n:6)=6,50000000000000<br>'on''sus' mnRank(n:8)=38,7500000000000<br>''off''tr' mnRank(n:15)=31,2000000000000<br>''off''sus' mnRank(n:19)=18,8947368421053 | "On transient" vs "On sustained" p: 0,000117355544889015<br>"On transient" vs "Off transient" p: 0,00148185948014506<br>"On transient" vs "Off sustained" p: 0,232028632846844<br>"On sustained" vs "Off transient" p: 0,606545019399266<br>"On sustained" vs "Off sustained" p: 0,00425963676826905<br>"Off transient" vs "Off sustained" p: 0,0533107400328478 |
| Sup3 | C | activity correl between c | unpaired t-test | Som-Ctrl n=7 vs Som-Rptor KO n=5 | t(df:10) = -2,4019 | 0,037196 | | | |
| 4 | B | Trials/min | paired t-test | Lea :Som-Ctrl n=6 vs Som-Ctrl n=6 | t(df:5) = -5,0053 | 0,0040862 | | | |
| 4 | B | Trials/min | paired t-test | Lea :Som-Rptor KO n=5 vs Som-Rptor KO n=5 | t(df:4) = -2,608 | 0,059543 | | | |
| 4 | B | Trials/min | Signed Wilcoxon rank test | Relea :Som-Ctrl n=6 vs Som-Ctrl n=6 | z-val(ranksum:11) = 0,10483 | 0,91651 | | | |
| 4 | B | Trials/min | paired t-test | Relea :Som-Rptor KO n=5 vs Som-Rptor KO n=5 | t(df:4)=1,034 | 0,35953 | | | |
| 4 | B | Trials/min | unpaired t-test | Lea :Som-Ctrl n=6 vs Som-Rptor KO n=5 | t(df:9) = 0,9069 | 0,3881 | | | |
| 4 | B | Trials/min | unpaired t-test | Relea :Som-Ctrl n=6 vs Som-Rptor KO n=5 | t(df:9)=1,3676 | 0,20462 | | | |
| 4 | C | Speed | paired t-test | Lea :Som-Ctrl n=6 vs Som-Ctrl n=6 | t(df:5)=-6,0773 | 0,0017433 | | | |
| 4 | C | Speed | paired t-test | Lea :Som-Rptor KO n=5 vs Som-Rptor KO n=5 | t(df:4)=-3,1273 | 0,035277 | | | |
| 4 | C | Speed | paired t-test | Relea :Som-Ctrl n=6 vs Som-Ctrl n=6 | t(df:5)=-0,080815 | 0,93872 | | | |
| 4 | C | Speed | paired t-test | Relea :Som-Rptor KO n=5 vs Som-Rptor KO n=5 | t(df:4)=0,13121 | 0,90195 | | | |
| 4 | C | Speed | unpaired t-test | Lea :Som-Ctrl n=6 vs Som-Rptor KO n=5 | t(df:9)=-0,004818 | 0,99626 | | | |
| 4 | C | Speed | unpaired t-test | Relea :Som-Ctrl n=6 vs Som-Rptor KO n=5 | t(df:9)=0,55865 | 0,59003 | | | |
| 4 | D | % Success trials | paired t-test | Lea :Som-Ctrl n=6 vs Som-Ctrl n=6 | t(df:5)=-4,4234 | 0,0068699 | | | |
| 4 | D | % Success trials | Signed Wilcoxon rank test | Lea :Som-Rptor KO n=5 vs Som-Rptor KO n=5 | z-val(ranksum:3)=1,3416 | 0,17971 | | | |
| 4 | D | % Success trials | paired t-test | Relea :Som-Ctrl n=6 vs Som-Ctrl n=6 | t(df:5)=-3,3157 | 0,021108 | | | |
| 4 | D | % Success trials | Signed Wilcoxon rank test | Relea :Som-Rptor KO n=5 vs Som-Rptor KO n=5 | z-val(ranksum:0)=-1 | 0,31731 | | | |
| 4 | D | % Success trials | Wilcoxon Rank test | Lea :Som-Ctrl n=6 vs Som-Rptor KO n=5 | z-val(ranksum:51)=2,7765 | 0,0054941 | | | |
| 4 | D | % Success trials | Wilcoxon Rank test | Relea :Som-Ctrl n=6 vs Som-Rptor KO n=5 | z-val(ranksum:49)=2,3359 | 0,019497 | | | |
| 4 | E | % Lick in rwd zone | paired t-test | Lea :Som-Ctrl n=6 vs Som-Ctrl n=6 | t(df:5)=-8,8124 | 0,00031239 | | | |
| 4 | E | % Lick in rwd zone | paired t-test | Lea :Som-Rptor KO n=5 vs Som-Rptor KO n=5 | t(df:4)=0,88727 | 0,42506 | | | |
| 4 | E | % Lick in rwd zone | paired t-test | Relea :Som-Ctrl n=6 vs Som-Ctrl n=6 | t(df:5)=-1,5974 | 0,17106 | | | |
| 4 | E | % Lick in rwd zone | Signed Wilcoxon rank test | Relea :Som-Rptor KO n=5 vs Som-Rptor KO n=5 | z-val(ranksum:0)=-1 | 0,31731 | | | |
| 4 | E | % Lick in rwd zone | Wilcoxon Rank test | Lea :Som-Ctrl n=6 vs Som-Rptor KO n=5 | z-val(ranksum:51)=2,6717 | 0,0075462 | | | |
| 4 | E | % Lick in rwd zone | Wilcoxon Rank test | Relea :Som-Ctrl n=6 vs Som-Rptor KO n=5 | z-val(ranksum:51)=2,7096 | 0,0067359 | | | |
| 4 | F | % Accuracy | paired t-test | Lea :Som-Ctrl n=6 vs Som-Ctrl n=6 | t(df:5)=-4,0669 | 0,0096631 | | | |
| 4 | F | % Accuracy | Signed Wilcoxon rank test | Lea :Som-Rptor KO n=5 vs Som-Rptor KO n=5 | z-val(ranksum:3)=1,3416 | 0,17971 | | | |
| 4 | F | % Accuracy | paired t-test | Relea :Som-Ctrl n=6 vs Som-Ctrl n=6 | t(df:5)=-2,8834 | 0,034451 | | | |
| 4 | F | % Accuracy | Signed Wilcoxon rank test | Relea :Som-Rptor KO n=5 vs Som-Rptor KO n=5 | z-val(ranksum:0)=-1 | 0,31731 | | | |
| 4 | F | % Accuracy | Wilcoxon Rank test | Lea :Som-Ctrl n=6 vs Som-Rptor KO n=5 | z-val(ranksum:51)=2,7765 | 0,0054941 | | | |
| 4 | F | % Accuracy | Wilcoxon Rank test | Relea :Som-Ctrl n=6 vs Som-Rptor KO n=5 | z-val(ranksum:49)=2,3359 | 0,019497 | | | |
| 4 | G | Learning index | paired t-test | Lea :Som-Ctrl n=6 vs Som-Ctrl n=6 | t(df:5)=-9,338 | 0,00023719 | | | |
| 4 | G | Learning index | paired t-test | Lea :Som-Rptor KO n=5 vs Som-Rptor KO n=5 | t(df:4)=-1,5795 | 0,18937 | | | |
| 4 | G | Learning index | paired t-test | Relea :Som-Ctrl n=6 vs Som-Ctrl n=6 | t(df:5)=-3,6918 | 0,01412 | | | |
| 4 | G | Learning index | Signed Wilcoxon rank test | Relea :Som-Rptor KO n=5 vs Som-Rptor KO n=5 | z-val(ranksum:0)=-1 | 0,31731 | | | |
| 4 | G | Learning index | unpaired t-test | Lea :Som-Ctrl n=6 vs Som-Rptor KO n=5 | t(df:9)=4,1322 | 0,0025511 | | | |
| 4 | G | Learning index | Wilcoxon Rank test | Relea :Som-Ctrl n=6 vs Som-Rptor KO n=5 | z-val(ranksum:51)=2,6779 | 0,0074078 | | | |
| 4 | J | Place corr. | Signed Wilcoxon rank test | Lea :Som-Ctrl n=40 vs Som-Ctrl n=40 | z-val(ranksum:213)=-2,6479 | 0,0080985 | | | |
| 4 | J | Place corr. | paired t-test | Lea :Som-Rptor KO n=32 vs Som-Rptor KO n=32 | t(df:31)=-0,87027 | 0,39084 | | | |
| 4 | J | Place corr. | paired t-test | Relea :Som-Ctrl n=40 vs Som-Ctrl n=40 | t(df:39)=-3,7532 | 0,00056836 | | | |

| | | | | | | |
|---|---|---|---|---|---|---|
| 4 | J | Place corr. | paired t-test | Relea :Som-Rptor KO n=32 vs Som-Rptor KO n=32 | t(df:31)=-1,9133 | 0,064989 |
| 4 | J | Place corr. | Wilcoxon Rank test | Lea :Som-Ctrl n=40 vs Som-Rptor KO n=32 | z-val(ranksum:1739)=3,1561 | 0,001599 |
| 4 | J | Place corr. | Wilcoxon Rank test | Relea :Som-Ctrl n=40vs Som-Rptor KO n=32 | z-val(ranksum:1655)=2,2042 | 0,027513 |
| 4 | N | % modulated cells | paired t-test | lea_st Som-Ctrl n=6 vs lea_end Som-Ctrl n=6 | t(df:5)=-4,554 | 0,0060903 |
| 4 | N | % modulated cells | paired t-test | lea_end Som-Ctrl n=6 vs relea_st Som-Ctrl n=6 | t(df:5)=4,1077 | 0,009285 |
| 4 | N | % modulated cells | paired t-test | relea_st Som-Ctrl n=6 vs relea_end Som-Ctrl n=6 | t(df:5)=-2,3972 | 0,061832 |
| 4 | N | % modulated cells | paired t-test | lea_end Som-Ctrl n=6 vs relea_end Som-Ctrl n=6 | t(df:5)=1,7186 | 0,14632 |
| 4 | N | % modulated cells | paired t-test | lea_st Som-Rptor KO n=5 vs lea_end Som-Rptor KO n=5 | t(df:4)=-0,31494 | 0,76855 |
| 4 | N | % modulated cells | paired t-test | lea_end Som-Rptor KO n=5 vs relea_st Som-Rptor KO n=5 | t(df:4)=-0,70611 | 0,51907 |
| 4 | N | % modulated cells | paired t-test | relea_st Som-Rptor KO n=5 vs relea_end Som-Rptor KO n=5 | t(df:4)=-0,042969 | 0,96779 |
| 4 | N | % modulated cells | paired t-test | lea_end Som-Rptor KO n=5 vs relea_end Som-Rptor KO n=5 | t(df:4)=-1,2462 | 0,28069 |
| Sup4 | E | Speed corr. | Signed Rank test | Lea :Som-Ctrl n=40 vs Som-Ctrl n=40 | z-val(ranksum:163)=-1,8886 | 0,058946 |
| Sup4 | E | Speed corr. | paired t test | Lea :Som-Rptor KO n=32 vs Som-Rptor KO n=32 | t(df:31)=1,2138 | 0,23399 |
| Sup4 | E | Speed corr. | Signed Rank test | Relea :Som-Ctrl n=40 vs Som-Ctrl n=40 | z-val(ranksum:401)=-0,12097 | 0,90371 |
| Sup4 | E | Speed corr. | paired t test | Relea :Som-Rptor KO n=32 vs Som-Rptor KO n=32 | t(df:31)=-1,0823 | 0,28745 |
| Sup4 | E | Speed corr. | Wilcoxon Rank test | Lea :Som-Ctrl n=40 vs Som-Rptor KO n=32 | z-val(ranksum:1602)=1,6035 | 0,10881 |
| Sup4 | E | Speed corr. | Wilcoxon Rank test | Relea :Som-Ctrl n=40 vs Som-Rptor KO n=32 | z-val(ranksum:1448)=-0,13032 | 0,89631 |
| Sup4 | F | Acceleration corr. | paired t test | Lea :Som-Ctrl n=40 vs Som-Ctrl n=40 | t(df:31)=1,6413 | 0,11085 |
| Sup4 | F | Acceleration corr. | Signed Rank test | Lea :Som-Rptor KO n=32 vs Som-Rptor KO n=32 | z-val(ranksum:295)=0,57967 | 0,56214 |
| Sup4 | F | Acceleration corr. | Signed Rank test | Relea :Som-Ctrl n=40 vs Som-Ctrl n=40 | z-val(ranksum:303)=-1,4382 | 0,15037 |
| Sup4 | F | Acceleration corr. | Signed Rank test | Relea :Som-Rptor KO n=32 vs Som-Rptor KO n=32 | z-val(ranksum:285)=0,39268 | 0,69456 |
| Sup4 | F | Acceleration corr. | Wilcoxon Rank test | Lea :Som-Ctrl n=40 vs Som-Rptor KO n=32 | z-val(ranksum:1358)=-1,1502 | 0,25004 |
| Sup4 | F | Acceleration corr. | Wilcoxon Rank test | Relea :Som-Ctrl n=40 vs Som-Rptor KO n=32 | z-val(ranksum:1465)=0,050996 | 0,95933 |
| Sup4 | G | Deceleration corr. | Signed Rank test | Lea :Som-Ctrl n=40 vs Som-Ctrl n=40 | t(df:31)=-1,6155 | 0,11634 |
| Sup4 | G | Deceleration corr. | paired t test | Lea :Som-Rptor KO n=32 vs Som-Rptor KO n=32 | t(df:31)=-1,3951 | 0,1729 |
| Sup4 | G | Deceleration corr. | Signed Rank test | Relea :Som-Ctrl n=40 vs Som-Ctrl n=40 | z-val(ranksum:578)=2,2581 | 0,023937 |
| Sup4 | G | Deceleration corr. | paired t test | Relea :Som-Rptor KO n=32 vs Som-Rptor KO n=32 | t(df:31)=0,70483 | 0,48618 |
| Sup4 | G | Deceleration corr. | unpaired t-test | Lea :Som-Ctrl n=40 vs Som-Rptor KO n=32 | t(df:70)=0,19071 | 0,84931 |
| Sup4 | G | Deceleration corr. | unpaired t-test | Relea :Som-Ctrl n=40 vs Som-Rptor KO n=32 | t(df:70)=0,26923 | 0,78854 |
